# Predicting the evolution of Lassa Virus endemic area and population at risk over the next decades

**DOI:** 10.1101/2021.09.22.461380

**Authors:** Raphaëlle Klitting, Liana E. Kafetzopoulou, Wim Thiery, Gytis Dudas, Sophie Gryseels, Anjali Kotamarthi, Bram Vrancken, Karthik Gangavarapu, Mambu Momoh, John Demby Sandi, Augustine Goba, Foday Alhasan, Donald S. Grant, Robert F. Garry, Allison R. Smither, Mark Zeller, Matthias G. Pauthner, Michelle McGraw, Laura D. Hughes, Sophie Duraffour, Stephan Günther, Marc A. Suchard, Philippe Lemey, Kristian G. Andersen, Simon Dellicour

## Abstract

Lassa fever is listed among the diseases that pose the greatest risks to public health by the World Health Organization. This severe viral hemorrhagic fever is caused by Lassa virus, a zoonotic pathogen that repeatedly spills over to humans from its rodent reservoirs. It is currently not known how climate change, transformations in land use, and human population growth could affect the endemic area of this virus, currently limited to parts of West Africa. By exploring the environmental data associated with virus occurrence, we show how temperature, precipitation and the presence of pastures determine ecological suitability for virus circulation. We project that regions in Central and East Africa will likely become suitable for Lassa virus over the next decades and estimate that the total population living in areas suitable for Lassa virus may grow from about 100 million to 700 million by 2070. By analysing geotagged viral genomes, we find that in the event of Lassa virus being introduced into a new suitable region, its spread might remain spatially limited over the first decades. Our results highlight how the endemic area of Lassa virus may expand well beyond West Africa in the next decades due to human impact on the environment, putting hundreds of million more people at risk of infection.

## Introduction

Along with other viral infections that have gained prominence in recent years^1–3^, Lassa fever (Lassa) is listed by the World Health Organization (WHO) as one of the diseases that pose the greatest public health risk^4,5^. Lassa is a viral hemorrhagic fever with variable but generally high case fatality rates^6^ for which efficacious countermeasures are currently lacking^7,8^.

To date, Lassa cases have only been reported in West Africa with Guinea, Liberia, Nigeria and Sierra Leone containing most endemic hotspots^9^. Nigeria, in particular, has seen a significant increase in incidence in recent years, and confirmed more than a thousand cases in 2020^10^. Increasingly, neighbouring countries, including Benin, Ghana, Ivory Coast, Mali and Togo, have also been reporting infections^11–14^, suggesting that the true range of Lassa virus may span a sizable part of West Africa.

Lassa is caused by Lassa virus^15^, a member of the *Arenaviridae* family (genus *Mammarenavirus*). Most infections occur through exposure to the excreta of infected *Mastomys natalensis*. These rodents often live in close contact with human communities^16^ and are regarded as the primary reservoir for the virus^17^. Humans likely contribute little to virus transmission and are considered dead-end hosts, based on studies of rodent biology^17^, ecology^15,20^, transmission dynamics^21,22^, and viral genomes^23–25^. While the virus can only spread where its reservoir is present, the range of *M. natalensis* extends beyond that of Lassa virus, spanning most of sub-Saharan Africa^26,27^. The factors underlying this difference in range between the virus and its reservoir have led to long-standing questions about suitability and may be multifactorial: Lassa virus might exclusively circulate within one of the six *M. natalensis* phylogroups or subspecies, namely A-I^28^, which is only found in West Africa^29–31^; other viruses present in *M. natalensis* populations may prevent Lassa virus circulation through competition^32,33^; closely related mammarenaviruses inducing cross-reactive immunity^34–36^ may prevent Lassa virus infection; and finally, Lassa virus prevalence may be influenced by different environmental determinants than its reservoir.

Like the rest of the world, African countries will increasingly be affected by climate change, with warming temperatures and more extreme, yet rarer, precipitation^37–39^. These changes, combined with an increasing pressure on land resources due to a considerable projected human population expansion, are expected to result in important transformations of land use throughout Africa^40–42^. In this study, we modelled how the endemic range of Lassa virus may evolve in the next five decades in response to climate change, human population growth, and land use changes. To identify factors driving suitability for virus circulation, we analysed the environmental data associated with the occurrence of Lassa virus for a set of putative explanatory factors. We find that annual mean temperature, annual precipitation and the presence of pastures are the main factors determining ecological suitability for Lassa virus circulation. Using projections of climate, land use, and population up to 2070, we estimated future ecological suitability and show that within decades, the range suitable for Lassa virus may extend well beyond West Africa. Using population projections, we estimate that the extended part of the suitable range will be home to hundreds of millions of people. To identify if, in case of introduction into a new region, specific environmental factors could slow the spread of the virus or even halt it — as proposed for the Niger and Benue rivers^23,43^ — we analysed geotagged viral genomes and environmental data. We find no evidence that the environmental factors we investigated, not even main rivers, limit virus dispersal. We show however, that over the first decades following a successful introduction into a new region, unless the virus spreads significantly faster than in current endemic areas, its propagation could remain spatially limited. By combining ecological niche modelling with spatially-explicit phylogeography, our study showcases how climate and land use change may transform the future risk of Lassa in Africa.

## Results

### Temperature, precipitation and pastures/rangeland are the main determinants of ecological suitability for Lassa virus

*M. natalensis* plays a critical role in the circulation of Lassa virus: as the main virus reservoir, but also as the source of most human infections^17,23,25^. Interestingly, while *M. natalensis* occupies a wide range spanning most sub-Saharan Africa^17^, Lassa virus has never been found outside of West Africa. This difference in range is poorly understood, but may in part be explained by the virus being more sensitive to environmental factors than its reservoir, as in the case of Sin Nombre virus^44^. For this other rodent-borne virus, environmental conditions can impact the abundance of the host, driving the population density of the reservoir below the threshold needed for virus maintenance^44–46^. Hence, to investigate factors that may determine ecological suitability, a measure of how suitable environmental conditions are for Lassa virus and *M. natalensis*, we conducted separate analyses for the virus and its reservoir species. We explored the environmental data associated with the occurrence of the virus (or its reservoir species) and found that annual mean temperature, annual precipitation and pastures/rangeland land coverage are the main determinants of ecological suitability for Lassa virus, while for *M. natalensis*, precipitation seems to be the most important factor.

To identify factors that determine ecological suitability for Lassa virus and *M. natalensis*, we built ecological niche models, considering temperature, precipitation, seven types of land cover and human population as potential determinants. Using a boosted regression trees^47^ (BRT) method, we searched for associations between known occurrences of the virus and its reservoir and the environmental conditions at those sites. As inputs for our models, we used occurrence records collated from online databases and the literature and environmental data obtained from the Inter-Sectoral Impact Model Intercomparison Project phase 2b (ISIMIP2b)^48^. To assess how each factor contributed to our models, we calculated their relative importance (RI). In the case of BRT models, RI is evaluated based on the number of times the factor is selected for splitting a tree, weighted by the squared improvement to the model as a result of each split, averaged over all trees^47^.

We found that for Lassa virus, three main factors contributed to the models: temperature (RI = 20.7%), precipitation (RI = 24.5%), pastures and rangeland land coverage (RI = 25.3%; **Fig. 1**). For *M. natalensis*, we found that precipitation was the main contributor (RI = 50.4%; **Fig. 1**). These findings suggest that temperature, precipitation and the presence of pastures/rangeland may be the main factors influencing ecological suitability for Lassa virus, but not its reservoir species, *M. natalensis*, for which only precipitation appears to be critical.

**Figure 1.**
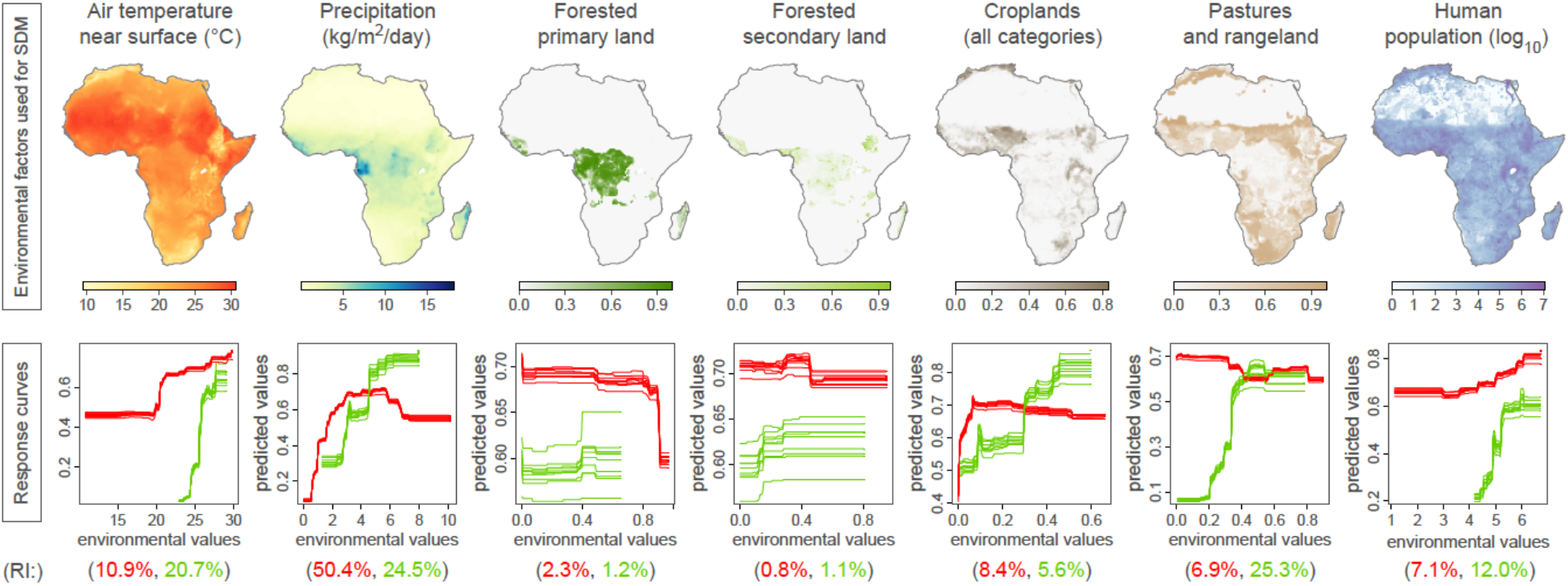
Environmental factors included in the ecological niche modelling (ENM) analyses of *Mastomys natalensis* and Lassa virus, and their corresponding ENM response curves. Response curves and relative importance (RI) obtained for the ENM analyses of *M. natalensis* and Lassa virus are coloured in red and green, respectively. The ten response curves reported for each ENM analysis correspond to ten independent boosted regression tree (BRT) repetitions. These response curves indicate the relationship between the environmental values and the response (i.e. the ecological suitability of *M. natalensis* or Lassa virus). In addition to the seven environmental factors displayed in this figure, two additional factors were also included in the ENM analyses, the non-forested primary land and non-forested secondary land.

To assess the relationship between each of our environmental factors and ecological suitability, we plotted response curves, which show how ecological suitability varies with one specific factor, while all others are kept constant at their mean. Ecological suitability values vary between 0 (unsuitable conditions) and 1 (highly suitable conditions). We found that temperatures below 25°C or values of pastures and rangeland land coverage below 20% seem unsuitable for the virus (ecological suitability ∼0; **Fig. 1**) but still appear relatively suitable (ecological suitability >0.4) for its reservoir species. These results indicate that even if *M. natalensis* may be found in areas with mean daily temperatures below 25°C and limited pastures and rangeland land coverage, Lassa virus is not likely to be present.

### Ecological niche modeling projects a likely expansion of the range suitable for Lassa virus

Our ecological niche modelling analyses showed that temperature, precipitation, and pastures/rangeland coverage are the main factors influencing ecological suitability for Lassa virus circulation. Due to climate change and increasing human pressure on land resources caused by population growth, these variables are expected to change in the next decades^40–42^. With these expected transformations, the overall area suitable for Lassa virus — also called the ecological niche of the virus^49^ — will likely undergo substantial changes and expand. To investigate this, we used climate and land cover projections from the year 2030 to 2070 to estimate the future ecological suitability for the virus across Africa. We found that the ecological niche of Lassa virus will likely expand as new regions become suitable, notably in Central and East Africa.

We used our ecological niche models to identify the areas suitable for Lassa virus throughout Africa, based either on current or projected values of temperature, precipitation, land cover and human population from the ISIMIP2b^48^. We considered environmental values projected at three time points (2030, 2050 and 2070) according to three climate scenarios: representative concentration pathways (RCPs) 2.6, 6.0 and 8.5 — which describe the evolution of global warming depending on different trajectories of greenhouse gases atmospheric concentrations^50^. For the present-day situation, our ecological niche maps for Lassa virus and *M. natalensis* were in agreement with previous estimates^51,27^, showing Lassa virus suitability across West Africa, predominantly in current endemic countries of Guinea, Sierra Leone, Liberia, and Nigeria (**Fig. 2A**).

**Figure 2.**
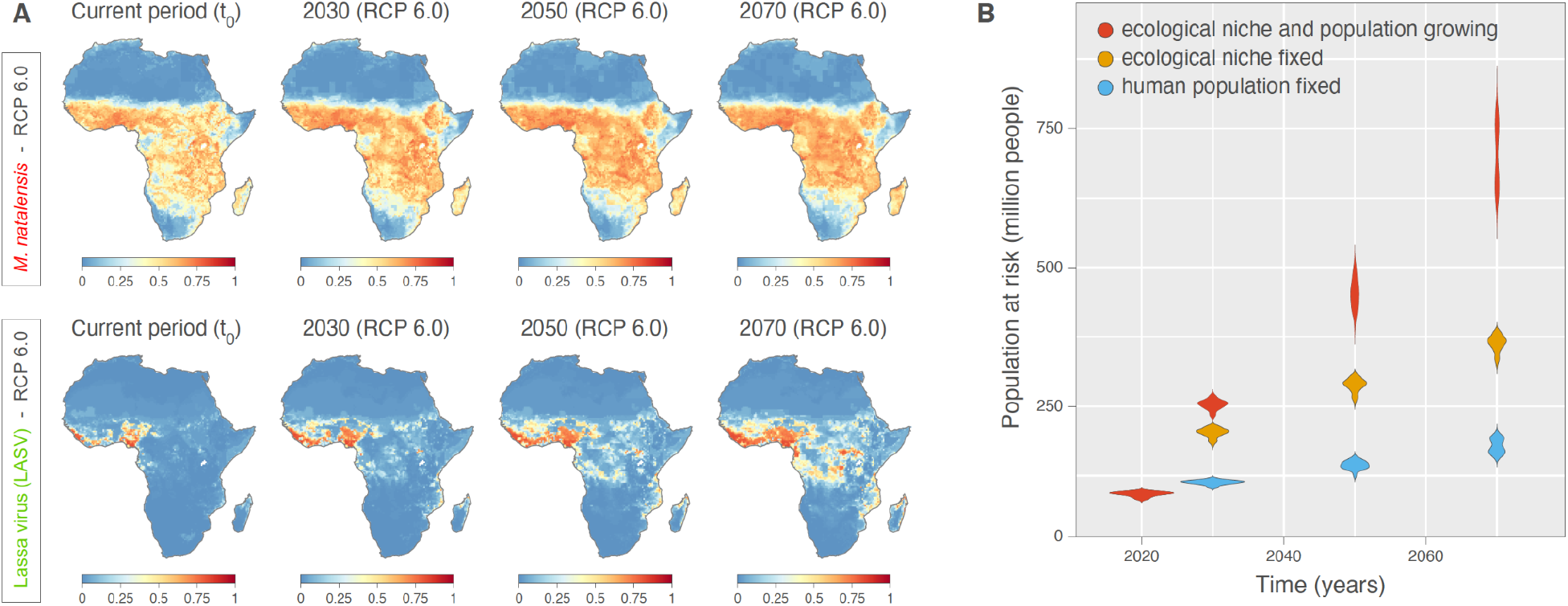
Projected ecological niche suitability of *Mastomys natalensis* and Lassa virus for the current period, 2030, 2050, and 2070 (A), as well as projections of the human population at risk of exposure to Lassa virus (B). Each future projection (i.e. for 2030, 2050, and 2070) was performed according to four different bias-adjusted global climate models and three different representative concentration pathways (RCPs), i.e. greenhouse gas concentration scenarios considered by the Intergovernmental Panel on Climate Change (IPCC): RCP 2.6, RCP 6.0, and RCP 8.5. Here, we only report the projections obtained under RCP 6.0 (see **Figure S1** for the other scenarios as well as **Figure S2** for explicit differences between current and future projections). For a specific time period, we report ecological niche suitability averaged over the projections obtained with the four different climatic models (see the text for further detail). For the estimations of the human population at risk of exposure to Lassa virus, we also re-estimate these projections while fixing the human population, i.e. not using the future projections of human population to estimate the number of people at risk (see also **Figure S3** for spatially-explicit estimation of future human exposure to Lassa virus and **Figure S6** for the estimations of the human population at risk of exposure to Lassa virus under all RCP scenarios).

At future time points, we found that the ecological niche of Lassa virus may substantially expand under both RCP 6.0 and RCP 8.5 (**Figs. 2A** and **S1**). RCP 2.6 and RCP 8.5 are the most extreme scenarios and refer to either stringent mitigation (RCP 2.6), or high-end emissions (RCP 8.5), while RCP 6.0 represents a medium-high emission scenario^48^. Focusing on RCP 6.0, we projected that by 2070, most of the region between Guinea and Nigeria will become suitable (ecological suitability >0.5) for Lassa virus (**Fig. 2A**). In addition, we found that several regions will likely become suitable in Central Africa, including in Cameroon and the Democratic Republic of the Congo (DRC), but also in East Africa, notably in Uganda. For *M. natalensis*, we found that irrespective of the scenario, the ecological niche will likely remain stable in range, with suitability values that increase over time and across the entire niche (**Figs. 2A** and **S1**). These results show that, considering a medium-high scenario of evolution of global warming (RCP 6.0), the ecological niche of Lassa virus may expand well beyond current endemic countries, notably into parts of Central and East Africa.

To investigate the factor(s) driving the expansion of the niche of Lassa virus, we represented, on a map of Africa, the environmental values for the main factors influencing ecological suitability at current and future time points (**Figs. S4** and **S5**). In Central and East Africa, areas showing an increased suitability for the virus under RCPs 6.0 and 8.5 also exhibited an increase in temperature and pastures/rangeland land coverage (**Figs. 2A, S4** and **S5**). Based on our observations, these two factors may thus primarily drive the expansion of the range suitable for Lassa virus.

### The population living where conditions are suitable for Lassa virus may increase dramatically by 2070, driven by a substantial population growth in both current and future areas suitable for the virus

Having found that the size of the ecological niche of Lassa virus will likely expand in the coming decades, we next investigated how this could affect the future number of people at risk of infection. To estimate the current and future human population in the Lassa virus niche, we considered population projections in areas with an estimated ecological suitability above 0.5 (**Fig. S3**). We focused again on three future time points (2030, 2050 and 2070) and three climate scenarios (RCPs 2.6, 6.0 and 8.5). We found that under RCP 6.0, the human population living in the niche of Lassa virus, where conditions are suitable for virus circulation, may increase from 92 million today (95% highest posterior density (HPD) interval: [83-98]) to 453 [414-498] million by 2050, and to 700 [624-779] million by 2070 (**Figs. 2B** and **S6, Table S1**).

This increase however, may be driven by demographic growth in current suitable areas rather than by the spatial expansion of the virus ecological niche^51^. To investigate this, we first examined current population numbers in areas suitable for Lassa virus in 2070 (scenario RCP 6.0) and found that they are currently home to 179 million people [159-199]. This result suggests that the population is expected to grow substantially throughout the entire niche of the virus (as projected in 2070), which will more than triple by 2070 (**Fig. 2B, Table S1**). When comparing the number of people that will live in current or future parts of the niche in 2070, we found that population growth should be comparable in both areas (**Table S1**). More specifically, our results show that by 2070, 363 [333-384] million people may be exposed to Lassa virus infection in current suitable areas and that expansion of the ecological niche of the virus may put 337 [260-405] million more people at risk of infection.

### Phylogeographic inference reveals that Lassa virus circulation is remarkably slow in endemic areas

In our ecological niche modelling analyses, we found that within a few decades, Lassa virus may circulate beyond its current endemic range in West Africa. If the virus is successfully introduced into new suitable regions, we estimated that hundreds of millions more people may be at risk of infection. The emergence of Ebola virus in West Africa and West Nile virus in North America show that zoonotic viruses can travel over long distances to effectively settle into new regions^52–55^. However, such events remain poorly understood and thus challenging to predict. While it is not possible to assess whether Lassa virus is likely to become established in a new environment, we can investigate how the virus may spread following a potential future introduction using, as *a priori* estimates, the parameters of virus dispersal in endemic areas. By analysing the spatiotemporal spread of the virus using geotagged viral genomes, we showed that Lassa virus dispersal in endemic areas is remarkably slow compared to other zoonotic viruses.

To infer the spatiotemporal spread of Lassa virus since the emergence of the four major clades^25^, we analysed publicly available genomic sequences associated with a sampling date and location using a spatially-explicit Bayesian phylogeographic approach^56^. The genome of Lassa virus is segmented into a large (L) and a small (S) segment that may reassort during coinfections in *M. natalensis*^57,58^. As reassortment may result in distinct evolutionary histories for the L and S segments, we analysed them in separate phylogeographic inferences. We also divided our analyses between four main clades (**Fig. S7**): The “MRU clade” groups the subclades circulating in the Mano River Union (MRU) and Mali (also called lineages IV and V); “NGA clade II”, “III” and “IV” correspond to the main clades circulating in Nigeria (also called lineages II, III and VI, respectively)^59^. The trees inferred by our phylogeographic analyses capture the spatiotemporal spread of the virus (**Fig. 3**), with each branch representing dispersal between an estimated start and end location, and associated with an estimated duration.

**Figure 3.**
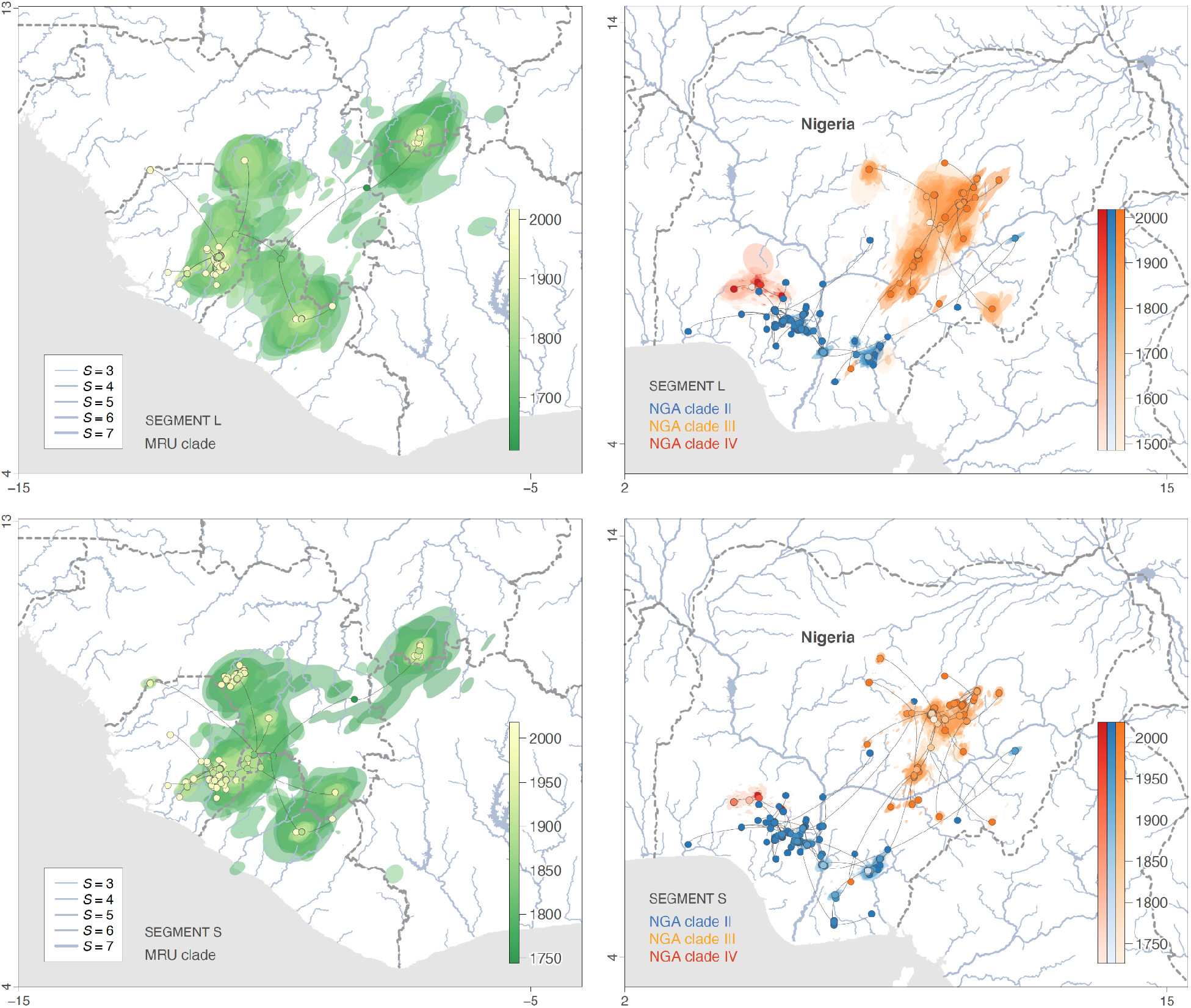
Spatiotemporal diffusion of Lassa virus lineages in the western Africa region and Nigeria. Maximum clade credibility (MCC) tree obtained by continuous phylogeographic inference based on 1,000 posterior trees. A separate phylogeographic analysis was performed for segments L and S as well as, in the case of the Nigerian data set, on clades II, III and IV. These MCC trees are superimposed on 80% HPDs reflecting phylogeographic uncertainty. Nodes of the trees, as well as HPD regions, are coloured according to their time of occurrence, and oldest nodes (and corresponding HPD regions) are here plotted on top of youngest nodes. The trees are superimposed on maps displaying the main rivers present in the study area and classified according to their Strahler number *S*, which measures the importance of a river by looking at the number of upstream rivers connected to it. International borders are represented by grey dashed lines. See also **Figures S9** and **S10** for visualisations clade by clade.

To investigate the spatial spread of the main clades, we represented the trees inferred by our phylogeographic analyses on maps, separately for the MRU and the Nigerian clades. We observed that in Nigeria, the main clades are confined to distinct areas: clades II and III circulates south and north of the Niger and Benue rivers, respectively, while clade IV is limited to states in the south west (Osun, Ekiti, Ondo, Kwara, **Fig. 3**). We also noted that sequences from the MRU clade grouped in three main clusters circulating respectively in eastern Sierra Leone, Guinea and Mali (**Fig. 3**). The strong geographic structure we observed in our phylogenetic trees was consistent across the S and L segments (**Figs. 3, S9**, and **S10**) and aligns with previous studies^23,24,43^. These findings suggest that, although the spread of Lassa virus encompasses hundreds of years (**Figs. 3, S9**, and **S10**), virus diversity is distinct across different areas.

To approximate how fast Lassa virus circulates in endemic areas (Manor River Union and Nigeria), we estimated the weighted lineage dispersal velocity^60^, which corresponds to the total distance covered by the dispersal events in our trees divided by the sum of their durations. We found that Lassa virus circulates with a posterior mean lineage dispersal velocity between 0.8 and 1 km/year (95% HPD interval for the S and L segments: [0.7 - 1.0] and [0.9 - 1.0]; **Table 1**). Our estimates of the weighted lineage dispersal velocity for each of the main clades show that virus circulation is slowest for the MRU clade and fastest for the Nigerian clade II (**Fig. S8**) but remains low overall (< 1.5 km/year). These results suggest that Lassa virus circulation is slow, which may in part explain why the main clades are confined to different areas within overall suitable regions (**Fig. 2**).

**Table 1.**
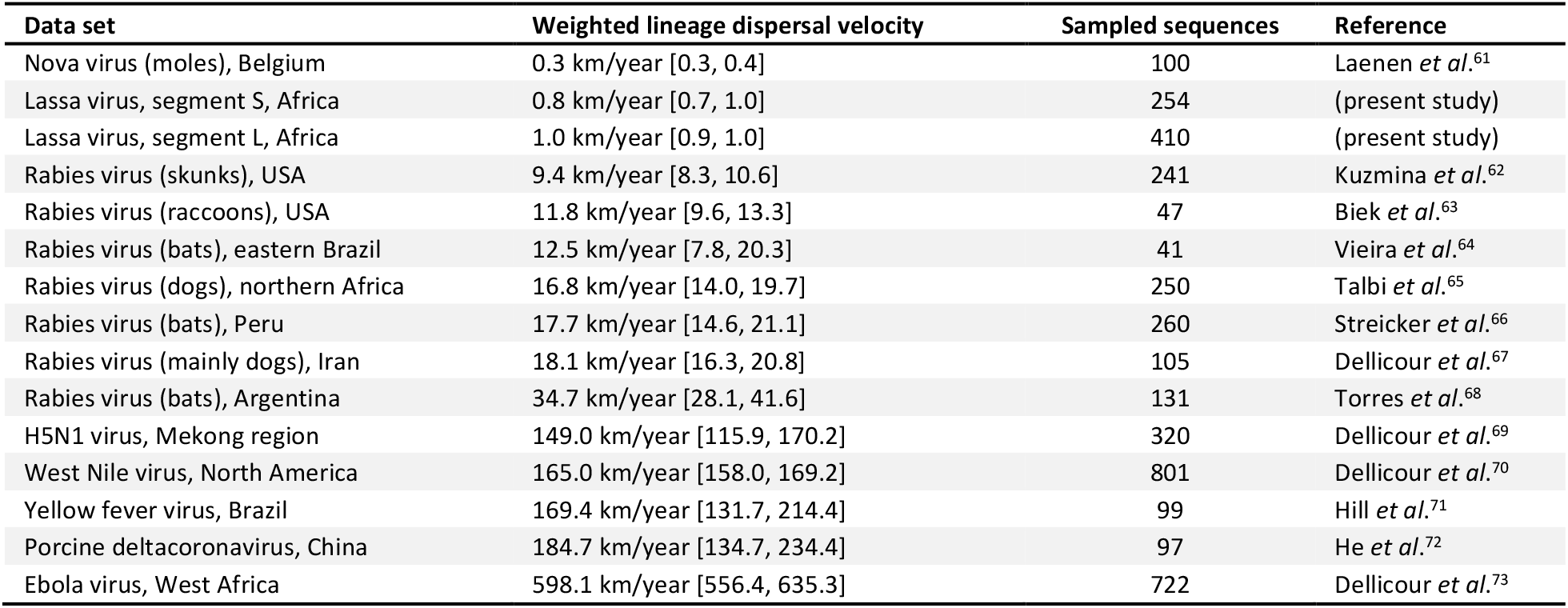
Comparison of lineage dispersal velocities estimated for different data sets. For each data set, we report both the posterior median estimate and the 95% HPD interval.

To determine how slow the velocity of Lassa virus circulation was compared to other zoonotic viruses, we assembled and sorted all published estimates of weighted lineage dispersal velocities (**Table 1**). We found that Lassa virus exhibits the slowest lineage dispersal velocity after Nova virus, while Ebola virus appeared to be the fastest. Our results indicate that Lassa virus circulation in endemic areas is particularly slow compared to other zoonotic viruses, possibly due to the small scale of the movements of its reservoir^17,26^.

### Lassa virus dispersal velocity and trajectory do not seem to be strongly affected by the environment

Our phylogeographic inferences show that Lassa virus circulation is remarkably slow in endemic areas, which may explain, at least in part, why the spatial spread of the main clades is limited, even within overall suitable regions. Nevertheless, since Lassa virus depends on *M. natalensis* for transmission, any environmental feature limiting the mobility of the reservoir may also impact virus dispersal. Main waterways in particular, have been proposed to act as barriers preventing the spread of Lassa virus, based on phylogenetic evidence that virus diversity is distinct across different sides of the Niger and Benue rivers in Nigeria^23,43^ (**Fig. 3**). Furthermore, for other viruses, such as rabies, there is evidence that environmental factors including elevation or croplands land coverage have an impact on virus lineage dispersal velocity^74,75^. To investigate how the environment may limit the propagation of Lassa virus following its potential future introduction into a new suitable region, we explored: (i) the impact of main waterways on the trajectory of Lassa virus dispersal, and (ii) the impact of nine different environmental factors on the velocity of Lassa virus circulation. We found that Lassa virus would likely spread unimpeded by environmental variables, and that major waterways may have limited impact on virus dispersal.

To determine if main waterways act as barriers to virus dispersal, we investigated whether Lassa virus tended to avoid crossing rivers based on our phylogeographic reconstructions. Using a least-cost path algorithm^76^, we computed the cost for the virus to travel through a landscape crossed by rivers based on both stream network data (**Table S6**) and the virus dispersal trajectory. We compared the cost of the observed spread inferred by our phylogeographic analyses to the cost computed under a null dispersal model that is unaware of rivers, and then estimated the statistical support for our test by approximating a Bayes Factor (BF) in favor of a cross-avoiding behavior. We repeated our test for a range of stream sizes considering different threshold values of the Strahler number (*S*) - a proxy for river stream size, based on a hierarchy of tributaries^77^. We found only moderate evidence (3 < BFs < 20)^78^ that the virus dispersal trajectory tends to avoid crossing rivers when considering rivers of intermediate sizes (*S* >4 and <5) and no evidence (BF <3) in the case of larger rivers (*S* >5, **Table S2**). Overall, our results provide no strong evidence that waterways may act as barriers to the dispersal of Lassa virus.

We next examined how environmental conditions may affect the velocity of Lassa virus circulation considering a set of nine environmental factors for which we collected geo-referenced data from public databases (**Fig. S11, Table S6**). For all virus dispersal events inferred by our phylogeographic analyses, we investigated whether the duration of the dispersal correlated with the environmental factors in our testing set. To assess these correlations, we computed an “environmental distance”, which corresponds to the distance of the dispersal event, weighted according to the environmental conditions along the path of dispersal. Our procedure only considers constant-in-time environmental values that do not reflect the climatic and land cover conditions during the earliest part of Lassa virus dispersal history, so we restricted our analyses to the most recent dispersal events (corresponding to tip branches of the trees from our phylogeographic reconstructions **Fig. S7**). We only found moderate evidence (3 < BFs < 20) that the presence of savannas may slow down viral circulation (**Tables S3 and S4**). These results suggest that the environmental factors considered in our analysis have no dramatic impact on the velocity of Lassa virus circulation.

### Simulations of Lassa virus spread show virus propagation may remain limited following introduction into a new suitable region

In the post-hoc analyses of our phylogeographic inferences, we did not identify any environmental factor that may prevent or notably slow down virus spread in a suitable environment. Hence, in case of introduction into a new suitable region, the main parameter that we can expect to limit Lassa virus propagation based on our analysis would be its slow lineage dispersal velocity. To illustrate how a slow lineage dispersal velocity may limit the spatial extent of virus spread following a potential introduction, we simulated virus dispersal based on the parameters inferred by our phylogeographic analyses (**Fig. 4**). We ran simulations over a 20-year period in two areas: one projected to become suitable for virus circulation by 2050 under scenario RCP 6.0 and the other one, under RCP 8.5. To simulate virus dispersal, we randomly sampled dispersal events inferred by our phylogeographic analyses for the Nigerian clade II, for which we have the largest number of sequences. We set the trajectory of dispersal events by selecting the ending location with a probability equal to the local ecological suitability (as projected in our ecological niche modelling analyses). By mapping the results of 1,000 simulations of virus dispersal, we show that Lassa virus would likely remain confined within a range of ∼200 km^2^ (**Fig. 4**), even when starting within a large suitable area (e.g. with scenario RCP 8.5, **Fig. 4 and S14**). Our simulations show how, if Lassa virus circulation is as slow as in current endemic areas, virus propagation would remain spatially limited over the first decades following its introduction into a new ecologically suitable area.

**Figure 4.**
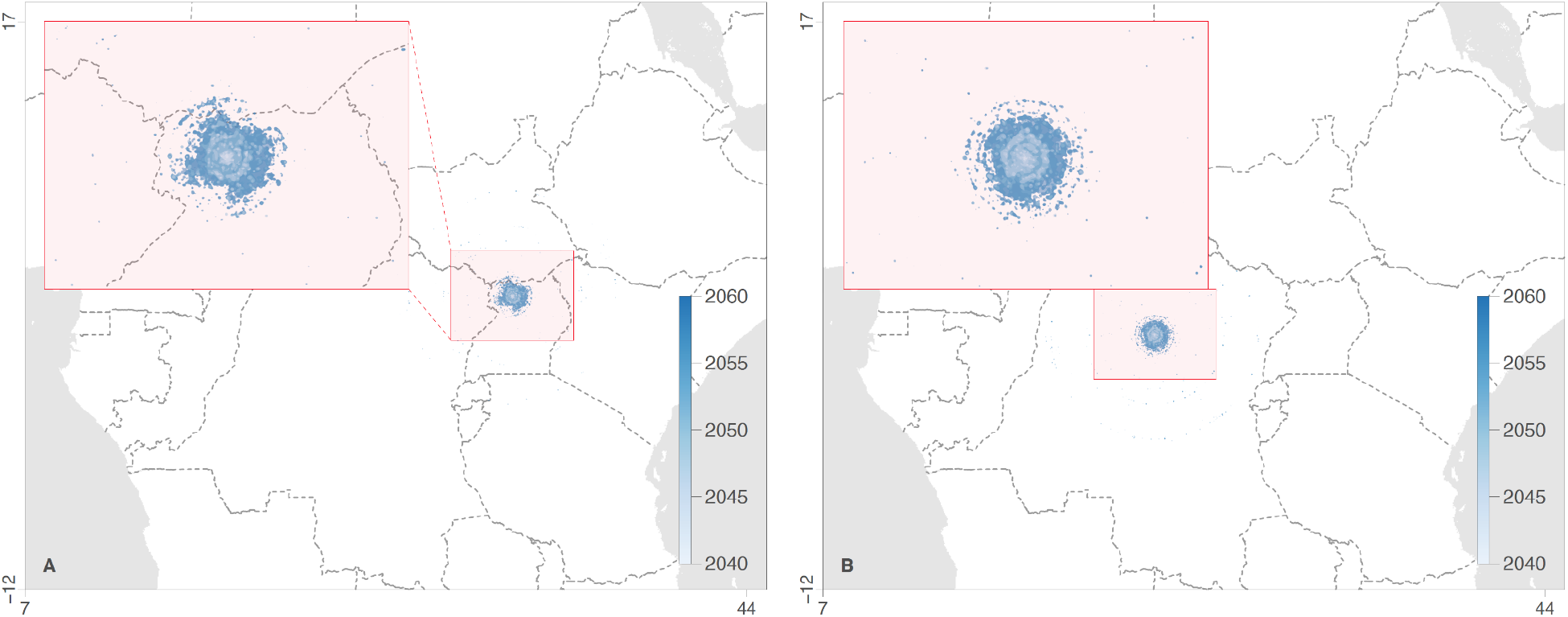
Phylogeographic simulations of viral spread following a successful introduction into a new ecologically suitable area. Phylogeographic simulations are based on the tree topologies inferred for Lassa virus clade II (segment S). Virus dispersal was constrained by ecological suitability using ecological niche projections for Lassa virus in 2050 according to scenarios RCP 6.0 (**A**) and RCP 8.5 (**B**). 95% HPD polygons are coloured according to time and based on 1,000 simulations starting from the same ancestral location. For each set of simulations, a zoom on the outcome is shown. For the illustration, five distinct phylogeographic simulations per scenario are also displayed in **Figure S14**.

## Discussion

Previous molecular dating studies have shown that Lassa virus has been circulating for at least 1,000 years and originated in present-day Nigeria, from where it spread to the West, reaching into the MRU region^11,25,79,80^. Lassa virus is considered endemic in Guinea, Liberia, Nigeria and Sierra Leone, but the virus likely circulates in other neighboring countries along its presumed dispersal path^11–14^. Although we do not attempt to precisely map the current range of Lassa virus, our ecological suitability estimates for the current period appear globally similar to the results obtained in an earlier study^27^ and show areas suitable for virus circulation across West Africa (see **Figure 2A**). Consistent with reports of Lassa virus infections in humans and rodents outside of endemic hotspots, our results suggest that the virus may be present in most coastal West African countries and Mali, prompting strengthened Lassa fever surveillance throughout the whole region.

*M. natalensis* is considered the primary reservoir of Lassa virus^16^ and it is still unclear why the distribution of this rodent species extends far beyond that of the virus, which is limited to West Africa^26^. Our analyses show that different environmental factors determine ecological suitability for the virus and its host, suggesting that the absence of Lassa virus beyond West Africa could be partly due to environmental constraints. To be able to estimate future ecological suitability for Lassa virus circulation, we have used projected environmental data with a low resolution (0.5 decimal degrees), due to the coarse scale of climate change projections^48,81^. This limited resolution reduced our ability to account for small-scale environmental variations that could affect suitability for Lassa virus; however, the good performance of our models (area under the receiver operating characteristic curves between 0.74 and 0.85) suggests that our approach provides a reasonable estimate of the distribution of Lassa virus infections.

In addition to environmental aspects, several other factors may also contribute to the difference in distribution between Lassa virus and its reservoir. As reported for other mammarenaviruses, the virus may only be present in the *M. natalensis* subtaxon A-I^30,31^. Records of Lassa virus infection in *M. natalensis* subtaxon A-II and in other rodent species including *M. erythroleucus* or *Hylomyscus pamfi*^82^ suggest, however, that susceptibility to Lassa virus infection may not be species or subtaxon specific. Other possible explanatory factors include intra-host competition between different viruses^32,83^, or cross-immunity due to the circulation of closely related viruses^34,35^. As most other old world arenaviruses that circulate in *M. natalensis* are found only in East Africa^30,31^, there is little data to assess these two mechanisms based on field data. Only in Mayo-Ranewo, eastern Nigeria, rodent trapping studies have identified a Mobala-like virus in *M. natalensis*^84^, which does not seem to effectively restrict the transmission of Lassa virus, as infections are often reported in that part of Nigeria^85,86^.

In our phylogeographic analyses we find that Lassa virus mostly spreads on a small spatial scale, with relatively few long-distance dispersal events (**Fig.3**) but we do not identify environmental factors that seem to strongly restrict or slow down virus spread. Using a phylogeographic simulation procedure, we also show that a slow lineage dispersal velocity would likely result in a limited spatial propagation if Lassa virus was successfully introduced in a new ecologically suitable area. The slow spread of Lassa virus is likely due to the small scale of the movements of its reservoir, as suggested by genetic studies showing that *M. natalensis* rodents travel rarely outside of their commensal habitat and are prone to high levels of consanguinity^26,17^. However, it is surprising that Lassa virus - and by implication its reservoir - seem unrestricted by the environment. Of note, our results were not always consistent between the S and L segments, possibly due to the lower number of L sequences (255) in our data set compared to S sequences (411). More generally, the number of genomic sequences in our datasets may offer limited power for the tests we used to assess the possible impact of rivers and other environmental factors on virus spread. Hence, a larger sampling of Lassa virus genomes throughout the virus range would allow for better evaluation of the role of the environment in limiting spread.

In our study, we use phylogeographic simulations to highlight how, in the absence of restrictions from the environment, a slow lineage dispersal velocity may limit the propagation of Lassa virus in case of introduction into a new ecologically suitable area. We use these simulations for illustration and not prediction as the dispersal dynamics upon virus emergence in a new region are unclear. The virus may spread swiftly through an immunologically naïve rodent population, but the low mobility of the rodent reservoir could still limit the velocity of virus dispersal on a larger scale. Many other elements may come into play, such as a possible change in reservoir host species, or the co-circulation of closely related viruses. Nevertheless, it is worth pointing out that following the emergence of Lassa virus in the MRU, virus circulation remained as slow - if not slower - as in Nigeria, as highlighted by our estimates of weighted lineage dispersal velocity (**Fig. S8**).

Our ecological niche analyses highlight a risk of expansion of Lassa virus towards regions in Central and East Africa that could be home to up to 337 million people by 2070 (**Table S1**). To reach the largest ecologically suitable regions we identify in DRC and Uganda, the virus would have to spread over several hundreds of kilometers and cross regions with low ecological suitability. Such long-distance movements likely allowed Lassa virus to reach the Mano River Union from Nigeria several hundred years ago^25^. This early part of the virus dispersal history, however, remains poorly understood and it is thus hard to predict if the virus is likely to travel across the African continent again. To provide a very conservative estimate of the future risk of exposure to Lassa virus, we can focus on population growth in the endemic range and leave aside its possible expansion. We estimated that population growth in endemic countries alone could alone put 341 [308-360] million people at risk of infection by 2070 (**Table S1**), compared to an estimated 92 [83-98] million today. A limitation to these estimates is that our population projections do not take into account migrations due to environmental and climate change pressures, which could affect projections in regions where extreme weather conditions are expected.

A large part of the population growth expected in endemic areas is driven by Nigeria (∼91%), a country that has reported an unusual increase in the number of reported Lassa fever cases over the last two years^85,86^. This uptick was not attributed to increased inter-human transmission^23,24^ or to the emergence of a specific viral strain^23,24^; but raised the question of a more intense circulation within the reservoir or of an improvement in surveillance and public awareness. To discriminate between these two hypotheses, we investigated the evolution of the overall genetic diversity of Lassa virus in the main Nigerian clade (clade II) over the past decades, using a coalescent approach that accounts for preferential sampling. We found that the effective population size of Nigeria clade II increased over the last years (segment S; **Fig. S13**), suggesting that the recent uptick in cases in Nigeria was not the sheer result of an improvement in surveillance. Hence, even if Lassa virus does not expand to new regions in the near future, the virus still actively circulates in increasingly populated endemic areas, and there is thus an urgent need for more efficient prophylactic and therapeutic countermeasures.

With anthropogenic climate change and an increasing impact of human activities on the environment, extensive studies of the ecology and spread of zoonotic and vector-borne diseases are needed to anticipate possible future changes in their distribution^87,88^. We showed that changes in temperature, precipitation and pastures/rangeland land coverage may expand the ecological niche of Lassa virus beyond current endemic areas, potentially exposing hundreds of million more people to Lassa. By simulating virus spread, we highlight that if virus propagation does not accelerate following introduction into new regions, the emerging circulation foci could remain limited to a small spatial scale over the first decades. Our study provides an example of how ecological niche modelling and spatially-explicit phylogeography can be effectively combined to investigate the future risk of a major zoonotic disease.

## Methods

### Ecological niche modelling of Mastomys natalensis and Lassa virus

We employed the boosted regression trees^89^ (BRT) approach implemented in the R package “dismo”^90^ to perform ecological niche modelling analyses of both Lassa virus and its host, *M. natalensis*. BRT is a machine learning method that allows to model complex non-linear relationships between the probability of occurrence and various predictor variables^90,91^. This approach aims to generate a collection of sequentially fitted regression trees that optimise the predictive probability of occurrence based on predictor values^89,91^, which can also be interpreted as a measure of ecological suitability. In a comprehensive review of distribution modeling methods, Elith *et al*.^89^ found BRT to perform best along with the maximum entropy method^92^. The BRT approach requires both presence and absence data. When unavailable, as this is the case for Lassa virus and its host, absence data can be approximated by random pseudo-absence points sampled from the study area (also referenced as the “background”). For Lassa virus, we only sampled pseudo-absences in raster cells in which the presence of *M. natalensis* has been recorded. This procedure avoids treating under-sampled areas as ecologically unsuitable for the virus, but also accounts for potential heterogeneity in sampling effort or surveillance^93,94^. Similarly, for *M. natalensis*, we only sampled pseudo-absences in raster cells in which the presence of at least one individual of another species belonging to the Muridae family has been recorded. Because it only requires a single occurrence record to consider a presence, we discarded all but one occurrence record per raster cell. We applied the same filtering step for the pseudo-absence points and simply discarded pseudo-absences falling in raster cells with occurrence data. To select the optimal number of trees in the BRT models, we used a spatial cross-validation procedure based on five spatially separated folds generated with the “blockCV” R package^95^. We employed a spatial rather than a standard cross-validation because the latter may overestimate the ability of the model to make reliable predictions when occurrence data are spatially auto-correlated^96^, which can frequently be the case. All BRT analyses were run and averaged over 10 cross-validated replicates, with a tree complexity set at 5, an initial number of trees set at 100, a learning rate of 0.005, and a step size of 10. We evaluated the inferences using the area under the receiver operating characteristic (ROC) curve, also simply referred to as “area under the curve” (AUC). Among replicates, AUC values ranged from 0.68 to 0.73 for *M. natalensis* (mean = 0.71), and from 0.74 to 0.85 for Lassa virus (mean = 0.79).

We obtained occurrence data for *M. natalensis* species from the Global Biodiversity Information Facility (http://www.gbif.org, accessed 2019-07-19), the Integrated Digitized Biocollections (https://www.idigbio.org, accessed 2020-01-04), the Field Museum of Natural History Zoological collections (https://collections-zoology.fieldmuseum.org, accessed 2019-12-13), and the African Mammalia database (http://projects.biodiversity.be/africanmammalia, accessed 2019-12-14). This data set was supplemented with the data available in the scientific literature (search for term “*Mastomys natalensis*”, in PubMed and Google). Duplicate records as well as records located in the ocean were excluded from the final data set, totalling 2,504 unique *M. natalensis* occurrence records. For 26 of those records, the location was not provided as spatial coordinates but as a locality (below or at the administrative level 4). Therefore, the latitude and longitude data correspond to that of the locality (determined as described in the subsection *Selection and preparation of viral sequences*; see below). Occurrence data for the *Muridae* family were obtained from the GBIF database. Duplicate records and records located in the ocean were excluded from the data set, totalling 10,806 unique *Muridae* occurrence records for the African continent. Occurrence data for Lassa virus were obtained by combining the data set from Fichet-Calvet & Rogers^97^ with records associated with sequences from our Lassa virus sequence data set (see below the subsection *Selection and preparation of viral sequences* for further detail), records of infected *M. natalensis* from our host occurrence data set and the data available in the scientific literature (search for term “Lassa virus” in PubMed). Duplicate records were discarded from the data set, resulting in 310 unique Lassa virus occurrence records. For two of those records, the location was not provided as spatial coordinates but as a locality (below or at the administrative level 4) so the latitude and longitude data corresponded to that of the locality (determined as described in the subsection *Selection and preparation of viral sequences*; see below). Our BRT models were trained on current environmental factors and then used to obtain estimates of future ecological niches for both Lassa virus and *M. natalensis*.

The BRT analyses were based on several environmental factors: harmonised present-day and future climate, land cover and population data available through the Inter-Sectoral Impact Model Intercomparison Project phase 2b (ISIMIP2b)^48^. The climate information consists of daily gridded near-surface air temperature and surface precipitation fields derived from four bias-adjusted^98^ global climate models (GCMs; GFDL-ESM2M^99^, HadGEM2-ES^100^, IPSL-CM5A-LR^101^, and MIROC5^102^) participating in the fifth phase of the Coupled Model Intercomparison Project (CMIP5^103^). We considered simulations conducted under historical climate forcings and RCPs 2.6, 6.0 and 8.5. In addition, we considered observed gridded temperature and precipitation from the concatenated products GSWP3 and EWEMBI^48^ for assessing the current (1986-2005) conditions. For land cover we use version 2 of the Land Use Harmonisation (LUH2^104^) providing historical and projected land cover states under a range of shared socioeconomic pathways (SSPs), and from which we consider SSP1-26, SSP4-6.0 and SSP8-85. Finally, we retrieve gridded population projections^105^ under SSP2-26. For each combination of product (GCM, GSWP3-EWEMBI LUH2, gridded population), scenario (historical, RCP, SSP) and analysis window (1986-2005, 2021-2040, 2041-2060, and 2061-2080), we compute the grid-scale temporal mean. For each scenario and time period, we estimated an index of human exposure (IHE) which corresponds to human population estimates (log^10^-transformed) in raster cells associated with an ecological suitability for Lassa virus above or equal to 0.5. Specifically, we used these IHE values to calculate the number of people at risk of exposure to Lassa virus. To investigate the specific effect of human population growth in current and future suitable areas, we also re-estimated future IHE values using (i) current population estimates with future projections of ecological suitability for Lassa virus to estimate population growth throughout current and future suitable areas, and (ii) future projections of human population with current projection of ecological suitability for Lassa virus to estimate the future population living in current suitable areas (Table S1). For each estimate, we calculated the mean and 95% HPD interval across all climatic models and ecological niche model replicates.

### Selection and curation of viral sequences

All publicly available sequences for Lassa virus were downloaded from the NCBI Nucleotide database (keywords: “lassa NOT mopeia NOT natalensis”). They were combined with recently generated sequences from Nigeria that have been sequenced as described previously by Kafetzopoulou and colleagues^106^ and that are publicly available on the website virological.org (https://virological.org/t/2019-lassa-virus-sequencing-in-nigeria-final-field-report-75-samples/291). We filtered the data by: i) excluding laboratory strains (adapted, passaged multiple times, recombinant, obtained from antiviral or vaccine experiments), ii) excluding sequences without a timestamp, (iii) keeping only sequences from a single timepoint (if multiple timepoints were available for a patient), iv) removing duplicates (when more than one sequence was available for a single strain), and v) excluding sequences from identified hospital epidemics or sequences for which the location corresponded to the site of hospitalisation. The remaining sequences were trimmed to their coding regions and arranged in sense orientation separately for the S segment (NP-NNN-GPC) and the L segment (L-NNN-Z). The sequences were aligned using MAFFT^107^ and inspected manually. At this step we discarded low quality sequences (manual curation) and very short sequences (combined ORF length <500nt). Since there is an overlap between the sequence data from the work of Siddle and colleagues^23^ and of Kafetzopoulou and colleagues^24^, we excluded sequences with zero or one mismatch between the two sets of sequences to ensure that there would not be duplicates in our data sets. Two types of alignments were generated. The alignments with all curated sequences regardless of the availability of detailed location information included 756 S segment sequences and 551 L segment sequences, respectively. The alignments with detailed location information included 411 S segment sequences and 255 L segment sequences, respectively. For the sequences with detailed location information, when no spatial coordinates were provided but only a name, spatial coordinates were determined using a combination of online platforms (Table S5). When several coordinates were available for one location, those matching across several data sets were kept, if the location was found in only one data set, the coordinates corresponding to the highest administrative level were kept.

### Inferring the dispersal history of Lassa virus lineages

We performed spatially-explicit phylogeographic reconstructions using the relaxed random walk (RRW) diffusion model^56^ implemented in BEAST 1.10^108^, which was coupled with the BEAGLE 3 library^109^ to improve computational performance. We modelled the nucleotide substitution process according to a GTR+Γ parameterisation^110^ and branch-specific evolutionary rates according to a relaxed molecular clock with an underlying log-normal distribution^111^. These phylogeographic analyses were based on the alignments of sequences associated with known spatial coordinates. For both the demographic and phylogeographic reconstructions, we ran a distinct BEAST analysis for each segment (L and S), sampling Markov chain Monte-Carlo (MCMC) chains every 10^5^ generations. We used Tracer 1.7^111^ for identifying the number of sampled trees to discard as burn-in, but also for inspecting the convergence and mixing, ensuring that estimated sampling size (ESS) values associated with estimated parameters were all >200. We used TreeAnnotator 1.10^112^ to obtain a maximum clade credibility (MCC) tree for each BEAST analysis. Finally, we used the R package “seraphim”^60^ to extract the spatiotemporal information embedded within trees obtained by spatially-explicit phylogeographic inference, as well as to estimate the weighted lineage dispersal velocity.

### Impact of environmental factors on the dispersal dynamics of Lassa virus lineages

Based on the spatially-explicit phylogeographic reconstructions, we performed two different kinds of analyses to investigate the impact of several environmental factors on the dispersal history and dynamics of Lassa virus lineages. First, we tested the impact of main rivers acting as potential barriers to Lassa virus dispersal (see Table S6 for the source of the original rivers shapefile). For this purpose, we used the least-cost path algorithm^76^ to compute the total cost for viral lineages to travel through a landscape crossed by rivers. This algorithm uses an underlying environmental raster to compute the minimum cost to move from one position to another. Here, we generated rasters by assigning a value of “1” to raster cells that were not crossed by a main river and a value of 1+*k* to raster cells crossed by a main river (raster resolution: ∼0.5 arcmin). Because the raster cells that were not crossed by a main river were assigned a uniform value of “1”, *k* thus defines the additional resistance to movement when the cell does contain such a potential landscape barrier^61,113^. In order to assess the impact of that rescaling parameter, we tested three different values for *k*: 10, 100 and 1000. Furthermore, as the notion of “main river” is arbitrary, we used different threshold values of the Strahler number *S* associated with each river to select the main rivers to consider in each analysis. In hydrology, *S* can be used as a proxy for stream size by measuring the branching complexity, i.e. the position of a river within the hierarchical river network. In practice, we compared the total cost computed for posterior trees with the total cost computed on the same trees along which we simulated a stochastic diffusion process under a null dispersal model^73^. Hereafter referred to as “simulated trees”, these trees were obtained by simulating a relaxed random walk process along the branches of trees sampled from the posterior distribution obtained by spatially-explicit phylogeographic inference^73^. Because this stochastic diffusion process did not take the position of rivers into account, we can expect the total cost to be lower for inferred trees under the assumption that viral lineages did tend to avoid crossing rivers. For each inferred or simulated tree, we computed the total cost *TC*, i.e. the sum of the least-cost values computed for each phylogenetic branch considered separately. Each “inferred” *TC* value (*TC*_inferred_) was then compared to its corresponding “simulated” value (*TC*_simulated_) by approximating a Bayes factor (BF) support as follows: BF = [*p*_*e*_/(1-*p*_*e*_)]/[0.5/(1-0.5)], where *p*_*e*_ is the posterior probability that *TC*_simulated_ > *TC*_inferred_, i.e. the frequency at which *TC*_simulated_ > *TC*_inferred_ in the samples from the posterior distribution. The prior odds is 1 because we can assume an equal prior expectation for *TC*_inferred_ and *TC*_simulated_.

Next, we tested the impact of several environmental variables, again described as rasters, on the dispersal velocity of Lassa virus lineages (Table S6, Figure S11): main rivers (as defined by selecting rivers with a *S* value higher than 2, 3, 4, 5, and 6), forest areas, grasslands, savannas, croplands, annual mean temperature, annual precipitation, and human population density. Except for the generated river rasters (see above), all these rasters had a resolution of ∼2.5 arcmin. For one-dimensional landscape features such as rivers, we had to resort to higher resolution rasters (∼0.5 arcmin) to obtain sufficiently precise pixelations (rasterizations) for this non-continuous environmental factor. Raster cells assigned to rivers would otherwise be exceptionally large given the size of the study area, which could potentially lead to artifactual results. Environmental rasters were tested as potential conductance factors (i.e. facilitating movement) as well as potential resistance factors (i.e. impeding movement). For each environmental variable, we also generated several distinct rasters with the following formula: *v*_*t*_ = 1 + *k*\*(*v*_*o*_/*v*_*max*_), where *v*_*t*_ is the transformed cell value, *v*_*o*_ the original cell value, and *v*_*max*_ the maximum cell value recorded in the raster. The rescaling parameter *k* here allows the definition and testing of different strengths of raster cell conductance or resistance, relative to the conductance/resistance of a cell with a minimum value set to “1”_70_. For each of the three environmental factors, we again tested three different values for *k* (i.e. 10, 100 and 1000). The following procedure can be summarised in three successive steps^114^: (i) based on environmental rasters, we computed environmental distance for each branch in inferred and simulated trees. These distances were computed using two different algorithms: the least-cost path and Circuitscape algorithm, the latter using circuit theory to accommodate uncertainty in the route taken^115^. For computational tractability, high resolution river rasters were only tested with the least-cost path algorithm. (ii) We estimated the correlation between time durations and environmental distances associated with each phylogenetic branch. Specifically, we estimated the statistic *Q* defined as the difference between the coefficient of determination obtained when branch durations are regressed against environmental distances computed on the environmental raster, and the coefficient of determination obtained when branch durations are regressed against environmental distances computed on a uniform “null” raster, i.e. a uniform raster with a value of “1” assigned to all its cells. We estimated *Q* for each tree and we thus obtained two distributions of *Q* values: one for inferred and one for simulated trees. We only considered an environmental raster as potentially explanatory if both its distribution of regression coefficients and its associated distribution of *Q* values were positive^116^. (iii) We evaluated the statistical support associated with a positive *Q* distribution (i.e. with at least 90% of positive values) by comparing it with its corresponding null distribution of *Q* values based on simulated trees. We formalised this comparison by approximating a BF support as defined above, but this time defining *p*_*e*_ as the posterior probability that *Q*_estimated_ > *Q*_simulated_, i.e. the frequency at which *Q*_estimated_ > *Q*_simulated_ in the samples from the posterior distribution^74^. For computational reasons, the “main rivers” rasters, which had to be associated with higher resolution (see above), were only tested as resistance factors with the least-cost-path algorithm.

### Phylogeographic simulations

We implemented a phylogeographic approach to simulate virus dispersal over a 20-year period following a successful introduction event within a new ecologically suitable area in 2050 under scenarios RCP 6.0 and RCP 8.5. We simulated viral lineage dispersal events by randomly sampling from the dispersal events inferred by our phylogeographic analyses. These simulations were performed under the assumption of no notable impact of underlying environmental factors. To set the trajectory of lineage dispersal events, we selected the ending location with a probability defined by the local ecological suitability. The starting point of those simulations was selected arbitrarily within the most suitable area of the extended part of the ecological niche estimated for LASV in 2050, and was thus different for simulations performed under scenarios RCP 6.0 and RCP 8.5.

### Inferring the demographic history of Lassa virus lineages

We performed demographic reconstructions using the flexible skygrid coalescent model^117^ implemented in BEAST 1.10^108^. The skygrid model allows to estimate the past evolution of the viral population effective size through time. For these analyses, we also modelled the nucleotide substitution process according to a GTR+Γ parameterisation^110^ and branch-specific evolutionary rates according to a relaxed molecular clock with an underlying log-normal distribution^111^. In the case of NGA clade II for which we inferred a recent increase in the global effective population size, we also performed a preferential sampling analysis^118^. By modeling the sampling times as a process dependent on effective population size, this complementary analysis allows to explicitly take into account heterogeneous sampling density through time, which can improve estimates of global effective population size^118^.

## Data Availability Statement

R scripts and related files needed to run all the ecological niche modelling and landscape phylogeographic analyses, as well as BEAST XML files, are all available at https://github.com/sdellicour/lassa_spreads.

## Acknowledgements

The authors thank David Pigott for sharing their environmental suitability predictions for *Mastomys natalensis*. Simon Dellicour is supported by the *Fonds National de la Recherche Scientifique* (FNRS, Belgium) and was previously funded by the *Fonds Wetenschappelijk Onderzoek* (FWO, Belgium). This research was supported by the National Institute Of Allergy And Infectious Diseases of the National Institutes of Health under Award Numbers U01AI151812, R01AI153044 and U19AI135995. The content is solely the responsibility of the authors and does not necessarily represent the official views of the National Institutes of Health. The research leading to these results has received funding from the European Research Council under the European Union’s Horizon 2020 research and innovation programme (grant agreement no. 725422-ReservoirDOCS), from the Wellcome Trust through project 206298/Z/17/Z (The Artic Network), and from the European Union’s Horizon 2020 project MOOD (grant agreement no. 874850). This research was supported by the Research and Innovation Programme of the European Union under H2020 grant agreement n°871029-EVA-GLOBAL. Philip Lemey acknowledges support by the Research Foundation - Flanders (*Fonds voor Wetenschappelijk Onderzoek* - *Vlaanderen*, G066215N, G0D5117N and G0B9317N).

## References

1. Morens, D. M. et al. The origin of COVID-19 and why it matters. Am. J. Trop. Med. Hyg. 103, 955–959 (2020).

2. Pierson, T. C. & Diamond, M. S. The emergence of Zika virus and its new clinical syndromes. Nature 560, 573–581 (2018).

3. Gates, B. The next epidemic - Lessons from Ebola. https://doi.org/10.1056/NEJMp1502918 (2015) doi:10.1056/NEJMp1502918.

4. World Health Organization. Lassa fever research and development (R&D) roadmap. https://www.who.int/publications/m/item/lassa-fever-research-and-development-(r-d)-roadmap (2018).

5. World Health Organization. Prioritizing diseases for research and development in emergency contexts. https://www.who.int/activities/prioritizing-diseases-for-research-and-development-in-emergency-contexts.

6. Akpede, G. O. et al. Caseload and case fatality of Lassa fever in Nigeria, 2001–2018: A specialist center’s experience and its implications. Front. Public Health 7, (2019).

7. Eberhardt, K. A. et al. Ribavirin for the treatment of Lassa fever: A systematic review and meta-analysis. Int. J. Infect. Dis. 87, 15–20 (2019).

8. Lukashevich, I. S., Paessler, S. & de la Torre, J. C. Lassa virus diversity and feasibility for universal prophylactic vaccine. F1000Res 8, (2019).

9. Bell-Kareem, A. R. & Smither, A. R. Epidemiology of Lassa fever. in 1–23 (Springer, 2021). doi:10.1007/82_2021_234.

10. Nigeria Centre for Disease Control. https://ncdc.gov.ng/diseases/sitreps/?cat=5&name=An%20update%20of%20Lassa%20fever%20outbreak%20in%20Nigeria.

11. Manning, J. T., Forrester, N. & Paessler, S. Lassa virus isolates from Mali and the Ivory Coast represent an emerging fifth lineage. Front. Microbiol. 6, (2015).

12. Dzotsi, E. K. et al. The first cases of Lassa fever in Ghana. Ghana. Med. J. 46, 166–170 (2012).

13. Patassi, A. A. et al. Emergence of Lassa fever disease in northern Togo: Report of two cases in Oti District in 2016. Case Rep. Infect. Dis. 2017, 8242313 (2017).

14. Yadouleton, A. et al. Lassa fever in Benin: Description of the 2014 and 2016 epidemics and genetic characterization of a new Lassa virus. Emerg. Microbes Infect. 9, 1761–1770 (2020).

15. McCormick, J. B. & Fisher-Hoch, S. P. Lassa fever. Curr. Top. Microbiol. Immunol. 262, 75–109 (2002).

16. Lecompte, E. et al. Mastomys natalensis and Lassa Fever, West Africa. Emerg. Infect. Dis. 12, 1971–1974 (2006).

17. Smither, A. R. & Bell-Kareem, A. R. Ecology of Lassa Virus. in 1–20 (Springer, 2021). doi:10.1007/82_2020_231.

18. Monath, T. P., Newhouse, V. F., Kemp, G. E., Setzer, H. W. & Cacciapuoti, A. Lassa virus isolation from Mastomys natalensis rodents during an epidemic in Sierra Leone. Science 185, 263–265 (1974).

19. Stephenson, E. H., Larson, E. W. & Dominik, J. W. Effect of environmental factors on aerosol-induced Lassa virus infection. J. Med. Virol. 14, 295–303 (1984).

20. Ogbu, O., Ajuluchukwu, E. & Uneke, C. J. Lassa fever in West African sub-region: An overview. J. Vector Borne Dis. 44, 1–11 (2007).

21. Ter Meulen, J. et al. Hunting of peridomestic rodents and consumption of their meat as possible risk factors for rodent-to-human transmission of Lassa virus in the Republic of Guinea. Am. J. Trop. Med. Hyg. 55, 661–666 (1996).

22. Lo Iacono, G. et al. Using modelling to disentangle the relative contributions of zoonotic and anthroponotic transmission: The case of Lassa fever. PLoS Negl. Trop. Dis. 9, (2015).

23. Siddle, K. J. et al. Genomic analysis of Lassa virus during an increase in cases in Nigeria in 2018. N. Engl. J. Med. 379, 1745–1753 (2018).

24. Kafetzopoulou, L. E. et al. Metagenomic sequencing at the epicenter of the Nigeria 2018 Lassa fever outbreak. Science 363, 74–77 (2019).

25. Andersen, K. G. et al. Clinical sequencing uncovers origins and evolution of Lassa virus. Cell 162, 738–750 (2015).

26. Lalis, A. & Wirth, T. Mice and men: An evolutionary history of Lassa fever. in Biodiversity and Evolution (eds. Grandcolas, P. & Maurel, M.-C.) 189–212 (Elsevier, 2018). doi:10.1016/B978-1-78548-277-9.50011-5.

27. Mylne, A. Q. N. et al. Mapping the zoonotic niche of Lassa fever in Africa. Trans. R. Soc. Trop. Med. Hyg. 109, 483–492 (2015).

28. Colangelo, P. et al. A mitochondrial phylogeographic scenario for the most widespread African rodent, Mastomys natalensis. Biol. J. Linn. Soc. 108, 901–916 (2013).

29. Colangelo, P. et al. A mitochondrial phylogeographic scenario for the most widespread African rodent, Mastomys natalensis. Biological Journal of the Linnean Society 108, 901–916 (2013).

30. Gryseels, S. et al. When viruses don’t go viral: The importance of host phylogeographic structure in the spatial spread of arenaviruses. PLoS Path. 13, e1006073 (2017).

31. Cuypers, L. N. et al. Three arenaviruses in three subspecific natal multimammate mouse taxa in Tanzania: Same host specificity, but different spatial genetic structure? Virus E vol. (2020) doi:10.1093/ve/veaa039.

32. Vazeille, M., Gaborit, P., Mousson, L., Girod, R. & Failloux, A.-B. Competitive advantage of a dengue 4 virus when co-infecting the mosquito Aedes aegypti with a dengue 1 virus. BMC Infect. Dis. 16, 318 (2016).

33. Chan, K. F. et al. Investigating viral interference between influenza A virus and human respiratory syncytial virus in a ferret model of infection. J. Infect. Dis. 218, 406–417 (2018).

34. Meunier, D. Y., McCormick, J. B., Georges, A. J., Georges, M. C. & Gonzalez, J. P. Comparison of Lassa, Mobala, and Ippy virus reactions by immunofluorescence test. Lancet 1, 873–874 (1985).

35. Howard, C. R. Antigenic diversity among the Arenaviruses. in The Arenaviridae (ed. Salvato, M. S.) 37–49 (Springer US, 1993). doi:10.1007/978-1-4615-3028-2_3.

36. Bhattacharyya, S., Gesteland, P. H., Korgenski, K., Bjørnstad, O. N. & Adler, F. R. Cross-immunity between strains explains the dynamical pattern of paramyxoviruses. Proc. Natl. Acad. Sci. U.S.A. 112, 13396–13400 (2015).

37. Coumou, D., Robinson, A. & Rahmstorf, S. Global increase in record-breaking monthly-mean temperatures. Climatic Change 118, 771–782 (2013).

38. Coumou, D. & Rahmstorf, S. A decade of weather extremes. Nat. Clim. Change 2, 491–496 (2012).

39. Bathiany, S., Dakos, V., Scheffer, M. & Lenton, T. M. Climate models predict increasing temperature variability in poor countries. Science Advances 4, eaar5809 (2018).

40. Arneth, A. Uncertain future for vegetation cover. Nature 524, 44–45 (2015).

41. Brandt, M. et al. Human population growth offsets climate-driven increase in woody vegetation in sub-Saharan Africa. Nat. Ecol. E vol. 1, 81 (2017).

42. Herrmann, S. M., Brandt, M., Rasmussen, K. & Fensholt, R. Accelerating land cover change in West Africa over four decades as population pressure increased. Com. Earth & Envir. 1, 1–10 (2020).

43. Ehichioya, D. U. et al. Phylogeography of Lassa virus in Nigeria. J. Virol. 93, e00929–19 (2019).

44. Luis, A. D., Douglass, R. J., Mills, J. N. & Bjørnstad, O. N. Environmental fluctuations lead to predictability in Sin Nombre hantavirus outbreaks. Ecology 96, 1691–1701 (2015).

45. Anderson, R. M., Jackson, H. C., May, R. M. & Smith, A. M. Population dynamics of fox rabies in Europe. Nature 289, 765–771 (1981).

46. Tian, H. et al. Anthropogenically driven environmental changes shift the ecological dynamics of hemorrhagic fever with renal syndrome. PLoS Pathogens 13, e1006198 (2017).

47. Elith, J., Leathwick, J. R. & Hastie, T. A working guide to boosted regression trees. J. Anim. Ecol. 77, 802–813 (2008).

48. Frieler, K. et al. Assessing the impacts of 1.5°C global warming – simulation protocol of the Inter-Sectoral Impact Model Intercomparison Project (ISIMIP2b). Geosci. Model Dev. 10, 4321–4345 (2017).

49. Soberón, J. & Nakamura, M. Niches and distributional areas: Concepts, methods, and assumptions. Proc. Natl. Acad. Sci. U.S.A. 106, 19644–19650 (2009).

50. Moss, R. H. et al. The next generation of scenarios for climate change research and assessment. Nature 463, 747–756 (2010).

51. Fichet-Calvet, E. & Rogers, D. J. Risk maps of Lassa fever in West Africa. PLoS. Negl. Trop. Dis. 3, (2009).

52. Lanciotti, R. S. et al. Origin of the West Nile virus responsible for an outbreak of encephalitis in the northeastern United States. Science 286, 2333–2337 (1999).

53. Alexander, K. A. et al. What factors might have led to the emergence of Ebola in West Africa? PLoS Negl. Trop. Dis. 9, e0003652 (2015).

54. Spengler, J., Ervin, E., Towner, J., Rollin, P. & Nichol, S. Perspectives on West Africa Ebola virus disease outbreak, 2013– 2016. Emerg. Infect. Dis. 22, 956 (2016).

55. Holmes, E. C., Dudas, G., Rambaut, A. & Andersen, K. G. The evolution of Ebola virus: Insights from the 2013–2016 epidemic. Nature 538, 193–200 (2016).

56. Lemey, P., Rambaut, A., Welch, J. J. & Suchard, M. A. Phylogeography takes a relaxed random walk in continuous space and time. Mol. Biol. Evol. 27, 1877–1885 (2010).

57. Lukashevich, I. S. Generation of reassortants between African arenaviruses. Virology 188, 600–605 (1992).

58. Vijaykrishna, D., Mukerji, R. & Smith, G. J. D. RNA virus reassortment: an evolutionary mechanism for host jumps and immune evasion. PLoS Path. 11, e1004902 (2015).

59. Whitmer, S. L. M. et al. New Lineage of Lassa Virus, Togo, 2016. Emerg. Infect. Dis. 24, 599 (2018).

60. Dellicour, S., Rose, R., Faria, N. R., Lemey, P. & Pybus, O. G. SERAPHIM: studying environmental rasters and phylogenetically informed movements. Bioinformatics 32, 3204–3206 (2016).

61. Laenen, L. et al. Spatio-temporal analysis of Nova virus, a divergent hantavirus circulating in the European mole in Belgium. Mol. Ecol. 25, 5994–6008 (2016).

62. Kuzmina, N. A. et al. The phylogeography and spatiotemporal spread of south-central skunk rabies virus. PLoS One 8, e82348 (2013).

63. Biek, R., Henderson, J. C., Waller, L. A., Rupprecht, C. E. & Real, L. A. A high-resolution genetic signature of demographic and spatial expansion in epizootic rabies virus. Proc. Natl. Acad. Sci. U.S.A. 104, 7993–7998 (2007).

64. Vieira, L. F. P., Pereira, S. R. F. G., Carnieli Jr, P., Tavares, L. C. B. & Kotait, I. Phylogeography of rabies virus isolated from herbivores and bats in the Espírito Santo State, Brazil. Virus Genes 46, 330–336 (2013).

65. Torres, C. et al. Phylodynamics of vampire bat-transmitted rabies in Argentina. Molecular Ecology 23, 2340–2352 (2014).

66. Streicker, D. G. et al. Host-pathogen evolutionary signatures reveal dynamics and future invasions of vampire bat rabies. Proc. Natl. Acad. Sci. U.S.A. 113, 10926–10931 (2016).

67. Dellicour, S. et al. Using phylogeographic approaches to analyse the dispersal history, velocity, and direction of viral lineages – application to rabies virus spread in Iran. Mol. Ecol. 28, 4335–4350 (2019).

68. Torres, C. et al. Phylodynamics of vampire bat-transmitted rabies in Argentina. Mol. Ecol. 23, 2340–2352 (2014).

69. Dellicour, S. et al. Incorporating heterogeneous sampling probabilities in continuous phylogeographic inference — Application to H5N1 spread in the Mekong region. Bioinformatics 36, 2098–2104 (2020).

70. Dellicour, S. et al. Phylogeographic and phylodynamic approaches to epidemiological hypothesis testing. bioRxiv (2020) doi:10.1101/788059.

71. Hill, S. C. et al. Genomic surveillance of yellow fever virus epizootic in São Paulo, Brazil, 2016 – 2018. PLoS Path. 16, e1008699 (2020).

72. He, W.-T. et al. Genomic epidemiology, evolution, and transmission dynamics of porcine deltacoronavirus. Mol. Biol. E vol. (2020).

73. Dellicour, S. et al. Phylodynamic assessment of intervention strategies for the West African Ebola virus outbreak. Nat. Commun. 9, 2222 (2018).

74. Dellicour, S. et al. Using viral gene sequences to compare and explain the heterogeneous spatial dynamics of virus epidemics. Mol. Biol. Evol. 34, 2563–2571 (2017).

75. Dellicour, S. et al. Epidemiological hypothesis testing using a phylogeographic and phylodynamic framework. Nat. Commun. 11, 5620 (2020).

76. Dijkstra, E. W. A note on two problems in connexion with graphs. Numer. Math. 1, 269–271 (1959).

77. Strahler, A. N. Quantitative analysis of watershed geomorphology. Eos, Transactions American Geophysical Union 38, 913–920 (1957).

78. Kass, R. E. & Raftery, A. E. Bayes factors. Journal of the American Statistical Association 90, 773–795 (1995).

79. Ehichioya, D. U. et al. Current molecular epidemiology of Lassa virus in Nigeria. J. Clin. Microbiol. 49, 1157 (2011).

80. Oloniniyi, O. K. et al. Genetic characterization of Lassa virus strains isolated from 2012 to 2016 in southeastern Nigeria. PLoS Negl. Trop. Dis. 12, e0006971 (2018).

81. Olesen, J. E. et al. Uncertainties in projected impacts of climate change on European agriculture and terrestrial ecosystems based on scenarios from regional climate models. Clim. Change 81, 123–143 (2007).

82. Olayemi, A. et al. New hosts of the Lassa virus. Sci. Rep. 6, 25280 (2016).

83. Zaidi, M. B. et al. Competitive suppression of dengue virus replication occurs in chikungunya and dengue co-infected Mexican infants. Parasit. Vectors 11, 378 (2018).

84. Olayemi, A. et al. Widespread arenavirus occurrence and seroprevalence in small mammals, Nigeria. Parasit. Vectors 11, 416 (2018).

85. ECHO Flash List. https://erccportal.jrc.ec.europa.eu/ECHO-Flash/ECHO-Flash-List/yy/2018/mm/2.

86. Nigeria Centre for Disease Control. https://ncdc.gov.ng/diseases/sitreps/?cat=5&name=An%20update%20of%20Lassa%20fever%20outbreak%20in%20Nigeria.

87. Pigott, D. M. et al. Local, national, and regional viral haemorrhagic fever pandemic potential in Africa: a multistage analysis. The Lancet 390, 2662–2672 (2017).

88. Kraemer, M. U. G. et al. Past and future spread of the arbovirus vectors Aedes aegypti and Aedes albopictus. Nature Microbiology (2019) doi:10.1038/s41564-019-0376-y.

89. Elith, J. et al. Novel methods improve prediction of species’ distributions from occurrence data. Ecography 29, 129–151 (2006).

90. Elith, J., Leathwick, J. R. & Hastie, T. A working guide to boosted regression trees. Journal of Animal Ecology 77, 802–813 (2008).

91. Dhingra, M. S. et al. Global mapping of highly pathogenic avian influenza H5N1 and H5Nx clade 2.3.4.4 viruses with spatial cross-validation. eLife 5, e19571 (2016).

92. Phillips, S. J., Anderson, R. P. & Schapire, R. E. Maximum entropy modeling of species geographic distributions. Ecological Modelling 190, 231–259 (2006).

93. Elith, J. et al. A statistical explanation of MaxEnt for ecologists. Diversity and Distributions 17, 43–57 (2011).

94. Phillips, S. J. et al. Sample selection bias and presence-only distribution models: Implications for background and pseudo-absence data. Ecological Applications 19, 181–197 (2009).

95. Valavi, R., Elith, J., Lahoz-Monfort, J. J. & Guillera-Arroita, G. blockCV: An r package for generating spatially or environmentally separated folds for k-fold cross-validation of species distribution models. Methods in Ecology and Evolution 10, 225–232 (2019).

96. Randin, C. F. et al. Are niche-based species distribution models transferable in space? J. Biogeogr. 33, 1689–1703 (2006).

97. Fichet-Calvet, E. & Rogers, D. J. Risk Maps of Lassa Fever in West Africa. PLoS Negl Trop Dis 3, (2009).

98. Lange, S. Bias correction of surface downwelling longwave and shortwave radiation for the EWEMBI dataset. Earth Syst. Dyn. 9, 627–645 (2018).

99. Dunne, J. P. et al. GFDL’s ESM2 global coupled climate–carbon earth system models. Part I: physical formulation and baseline simulation characteristics. J. Climate 25, 6646–6665 (2012).

100. Jones, C. D. et al. The HadGEM2-ES implementation of CMIP5 centennial simulations. Geosci. Model Dev. 4, 543–570 (2011).

101. Dufresne, J.-L. et al. Climate change projections using the IPSL-CM5 Earth System Model: from CMIP3 to CMIP5. Clim. Dyn. 40, 2123–2165 (2013).

102. Watanabe, M. et al. Improved climate simulation by MIROC5: Mean states, variability, and climate sensitivity. J. Climate 23, 6312–6335 (2010).

103. Taylor, K. E., Stouffer, R. J. & Meehl, G. A. An overview of CMIP5 and the experiment design. Bull. Amer. Meteor. Soc. 93, 485–498 (2012).

104. Hurtt, G. C. et al. Harmonization of global land-use change and management for the period 850-2100 (LUH2) for CMIP6. Geosci. Model Dev. 1–65 (2020) doi:https://doi.org/10.5194/gmd-2019-360.

105. Jones, B. & O’Neill, B. C. Spatially explicit global population scenarios consistent with the Shared Socioeconomic Pathways. Environ. Res. Lett. 11, 084003 (2016).

106. Kafetzopoulou, L. E. et al. Metagenomic sequencing at the epicenter of the Nigeria 2018 Lassa fever outbreak. Science 363, 74–77 (2019).

107. Katoh, K. & Standley, D. M. MAFFT Multiple Sequence Alignment Software version 7: Improvements in performance and usability. Mol. Biol. Evol. 30, 772–780 (2013).

108. Suchard, M. A. et al. Bayesian phylogenetic and phylodynamic data integration using BEAST 1.10. Virus Evol. 4, vey016 (2018).

109. Ayres, D. L. et al. BEAGLE 3: Improved performance, scaling, and usability for a high-performance computing library for statistical phylogenetics. Syst. Biol. (2019) doi:10.1093/sysbio/syz020.

110. Tavaré, S. Some probabilistic and statistical problems in the analysis of DNA sequences. Lectures Math. Life Sci. 17, 57–86 (1986).

111. Rambaut, A., Drummond, A. J., Xie, D., Baele, G. & Suchard, M. A. Posterior summarization in Bayesian phylogenetics using Tracer 1.7. Syst. Biol. 67, 901–904 (2018).

112. Suchard, M. A. et al. Bayesian phylogenetic and phylodynamic data integration using BEAST 1.10. Virus Evolution 4, vey016 (2018).

113. Dellicour, S. et al. Landscape genetic analyses of Cervus elaphus and Sus scrofa: comparative study and analytical developments. Heredity 123, 228–241 (2019).

114. Dellicour, S., Rose, R. & Pybus, O. G. Explaining the geographic spread of emerging epidemics: a framework for comparing viral phylogenies and environmental landscape data. BMC Bioinform. 17, 1–12 (2016).

115. McRae, B. H. Isolation by resistance. Evolution 60, 1551–1561 (2006).

116. Jacquot, M., Nomikou, K., Palmarini, M., Mertens, P. & Biek, R. Bluetongue virus spread in Europe is a consequence of climatic, landscape and vertebrate host factors as revealed by phylogeographic inference. Proc. R. Soc. Lond. B 284, 20170919 (2017).

117. Gill, M. S. et al. Improving Bayesian population dynamics inference: A coalescent-based model for multiple loci. Mol. Biol. Evol. 30, 713–724 (2013).

118. Karcher, M. D., Palacios, J. A., Bedford, T., Suchard, M. A. & Minin, V. N. Quantifying and mitigating the effect of preferential sampling on phylodynamic inference. PLoS Comput. Biol. 12, e1004789 (2016).

