## Supplementary tables and figures for "Predicting the evolution of Lassa Virus endemic area and population at risk over the next decades"

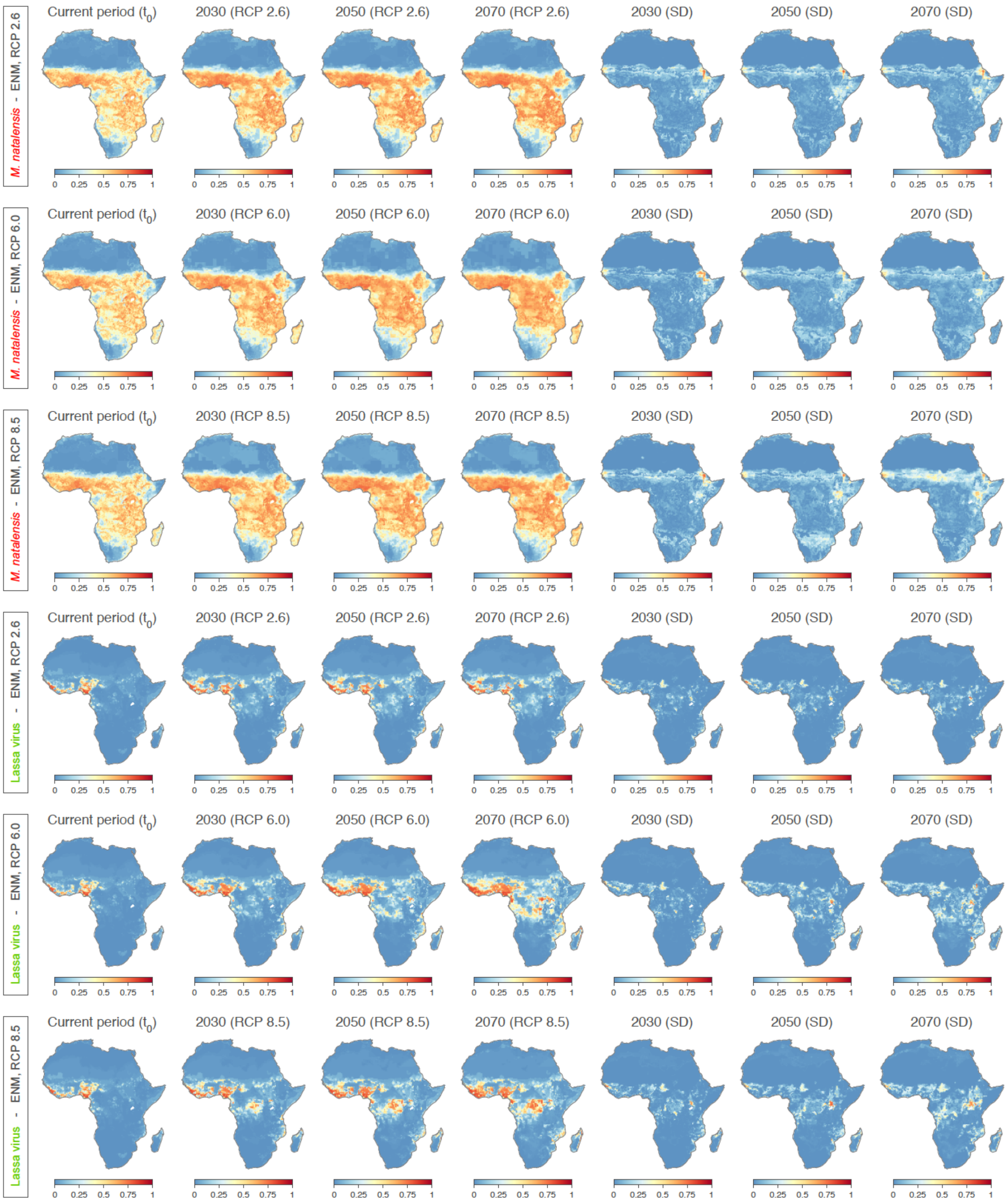


**Figure S1. Predicted ecological niche suitability for *Mastomys natalensis* and Lassa virus for the current period, 2030, 2050, and 2070, as well as associated standard deviation (SD) between climatic models.** Each future projection (i.e. for 2030, 2050, and 2070) was performed according to four different climatic models and three different representative concentration pathways (RCPs), i.e. greenhouse gas concentration scenarios defined by the Intergovernmental Panel on Climate Change (IPCC): RCP 2.6, RCP 6.0, and RCP 8.5. For a specific RCP and time period, we here report predicted probabilities averaged over the projections obtained with the four different climatic models (see the text for further detail). We also report the SD between the projections obtained for the four climatic models.

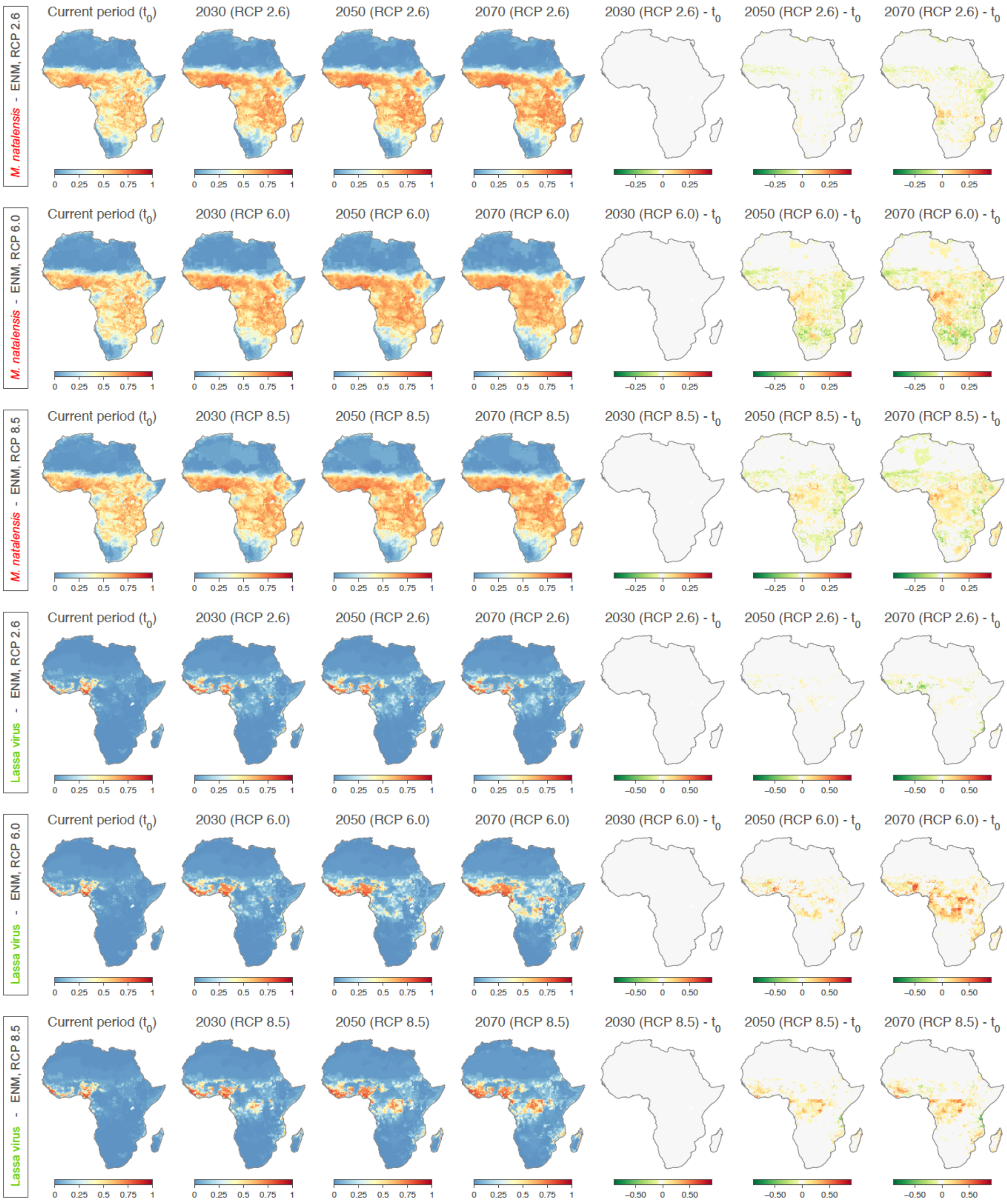


**Figure S2. Predicted ecological niche suitability for *Mastomys natalensis* and Lassa virus for the current period, 2030, 2050, and 2070, as well as differences between future and current projections.** Each future projection (i.e. for 2030, 2050, and 2070) was performed according to four different climatic models and three different representative concentration pathways (RCPs), i.e. greenhouse gas concentration scenarios defined by the Intergovernmental Panel on Climate Change (IPCC): RCP 2.6, RCP 6.0, and RCP 8.5 (IPCC scenario also commonly referred as ``business as usual''). For a specific RCP and time period, we here report predicted probabilities averaged over the projections obtained with the four different climatic models (see the text for further detail). We also report the difference between each future projection and the projection obtained for the current period $t_0$.

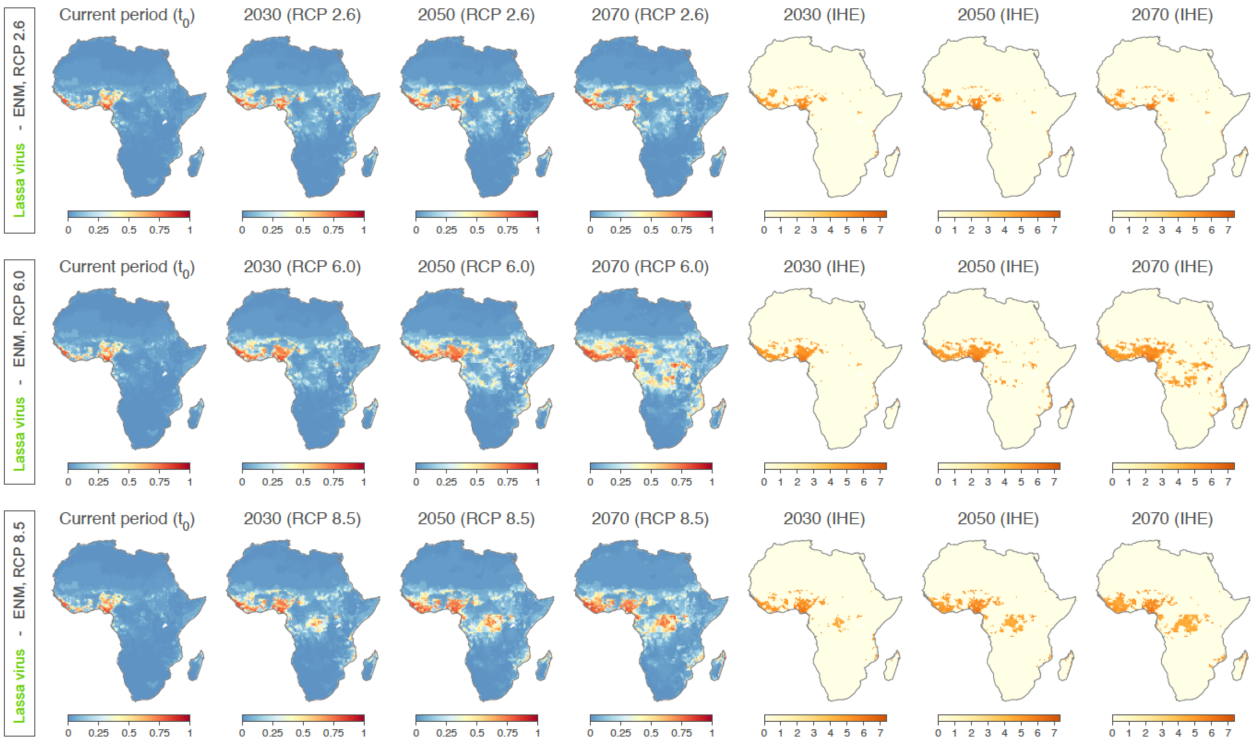


**Figure S3. Predicted ecological niche suitability of Lassa virus for the current period, 2030, 2050, and 2070, as well as associated estimations for an index of human exposure (IHE).** Each future projection (i.e. for 2030, 2050, and 2070) was performed according to three different representative concentration pathways (RCPs) and averaged over projections obtained according to four climatic models (see the text for further detail). IHE were obtained by only reporting human population estimates ($log_{10}$-transformed) associated with Lassa virus predicted probability higher than 0.5.

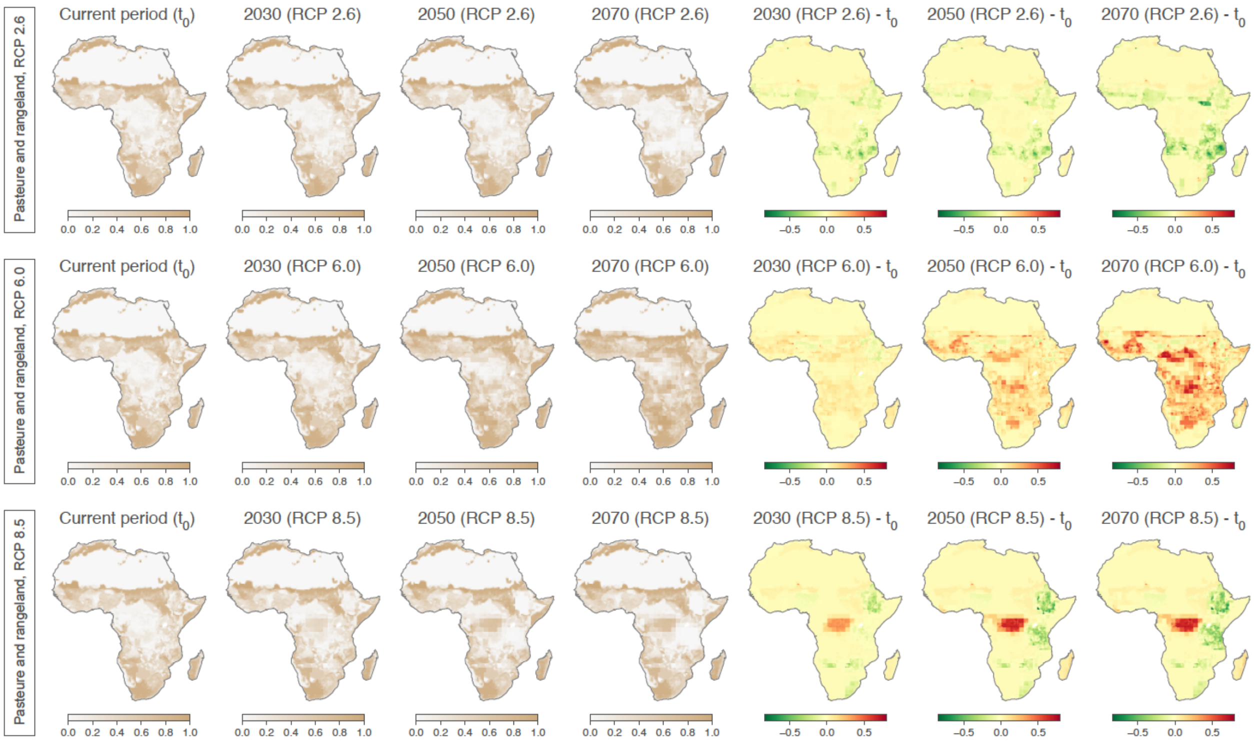


**Figure S4. Current and future estimates for the "pastures and rangeland" environmental factor used in the ENM analyses.** We here also report the difference between each future estimate and the estimate for the current period ($t_0$).

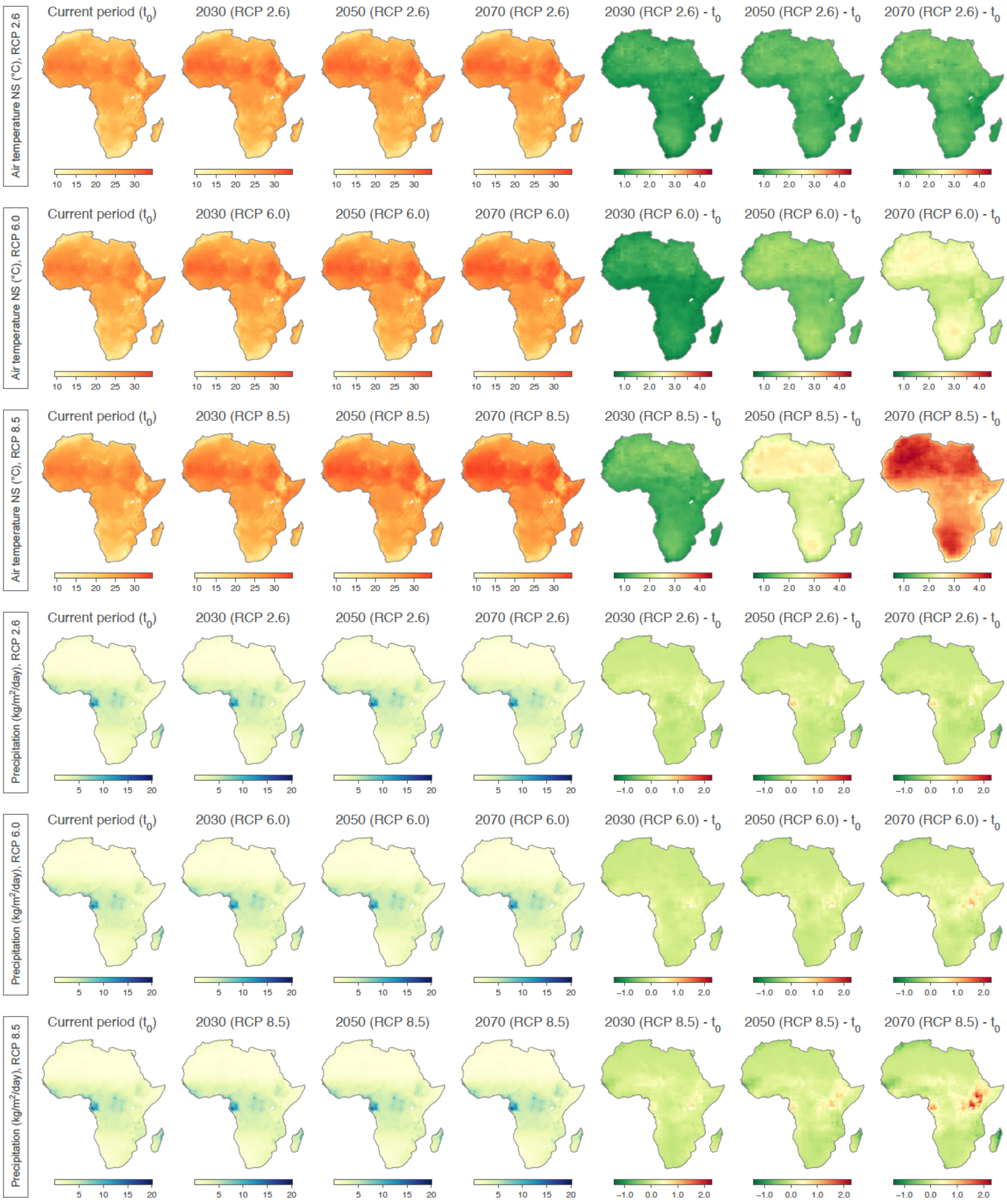


**Figure S5. Current and future estimates for the two climatic factors used in the ENM analyses: the air temperature near the surface (NS) and the precipitation rate.** We here also report the difference between each future estimate and the estimate for the current period ($t_0$).

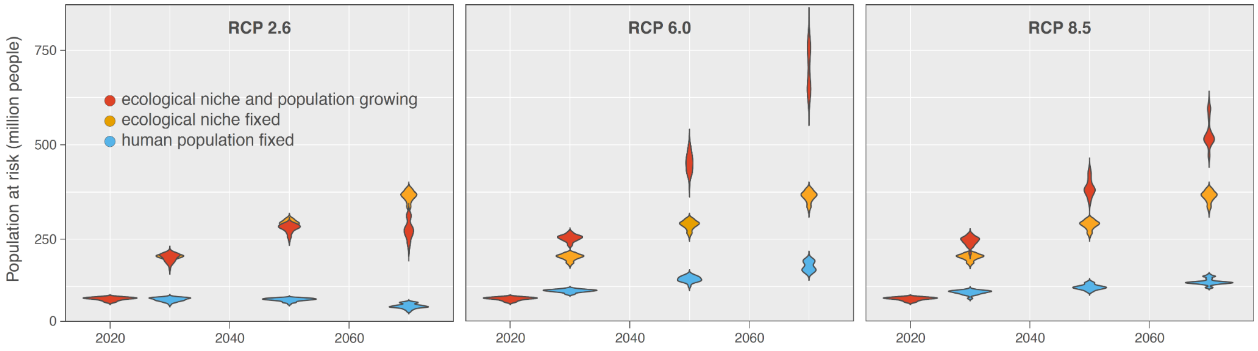


**Figure S6. Projections of the human population at risk of exposure to Lassa virus.** Each future projection (i.e. for 2030, 2050, and 2070) was obtained according to three different representative concentration pathways (RCPs) and averaged over projections obtained according to four climatic models (see the text for further detail). In addition, we also re-estimate these projections while fixing the human population, i.e. not using the future projections of human population to estimate the number of people at risk.

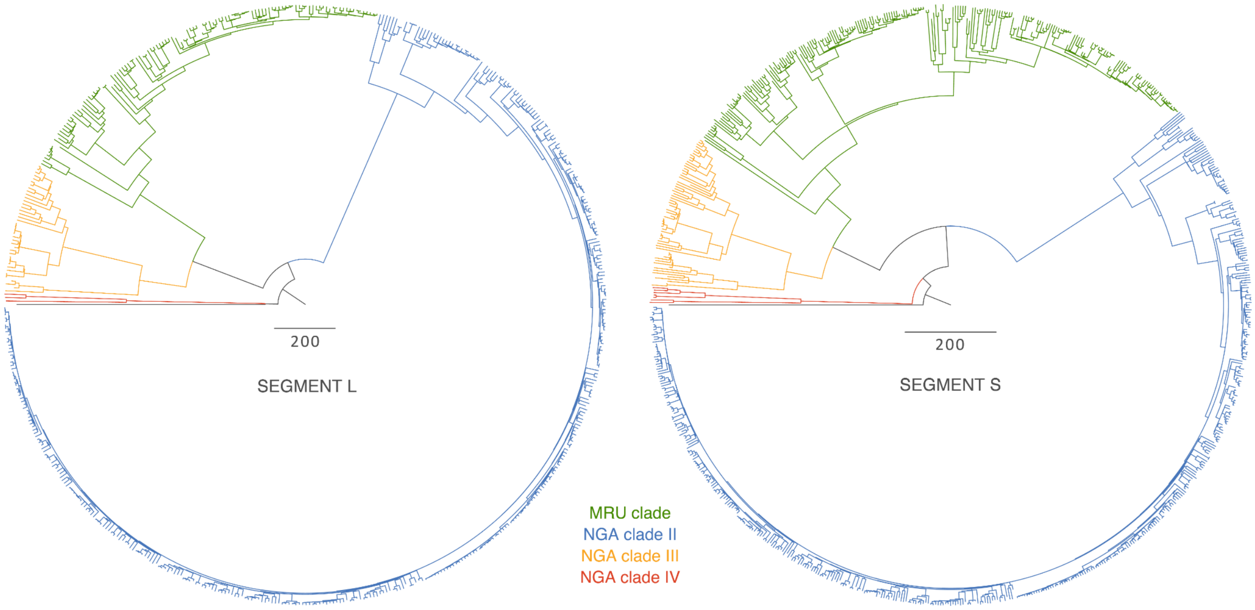


**Figure S7. Overall Lassa virus time-scaled phylogenetic trees based on segments L and S.** Clades are coloured according to the main LASV clades.

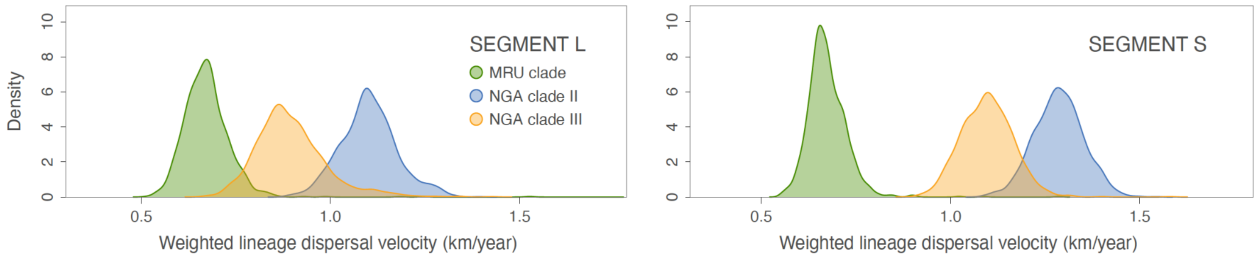


**Figure S8. Weighted dispersal velocities estimated for Lassa virus lineages.** These estimations are based on 1,000 posterior trees inferred by continuous phylogeographic inference.

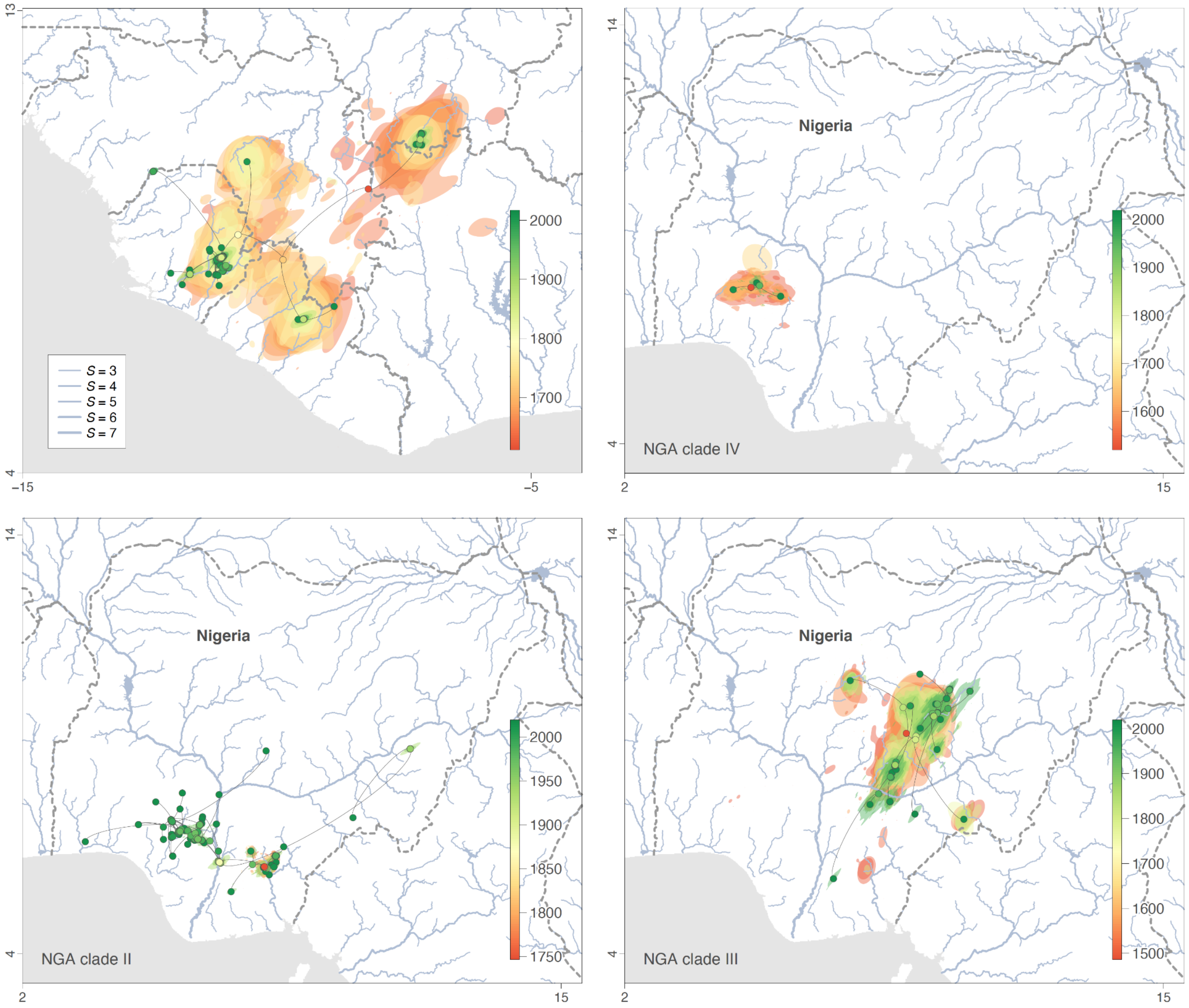


**Figure S9. Spatiotemporal diffusion of Lassa virus lineages in western Africa and Nigeria, based on the analysis of segment L.** MCC (maximum clade credibility) tree obtained by continuous phylogeographic inference based on 1,000 posterior trees. A distinct phylogeographic analysis has been based on segments L and S as well as, in the case of the Nigerian data set, on clades I, II and III. These MCC trees are superimposed on 80% HPD reflecting phylogeographic uncertainty. Nodes of the trees, as well as HPD regions, are coloured from red (the time to the most recent common ancestor, TMRCA) to green (most recent sampling time), and oldest nodes (and corresponding HPD regions) are here plotted on top of youngest nodes. The trees are superimposed on maps displaying the main rivers present in the study area and classified according to their Strahler number *S*, which measures the importance of a river by counting the number of upstream rivers connected to it. International borders are represented by white dashed lines.

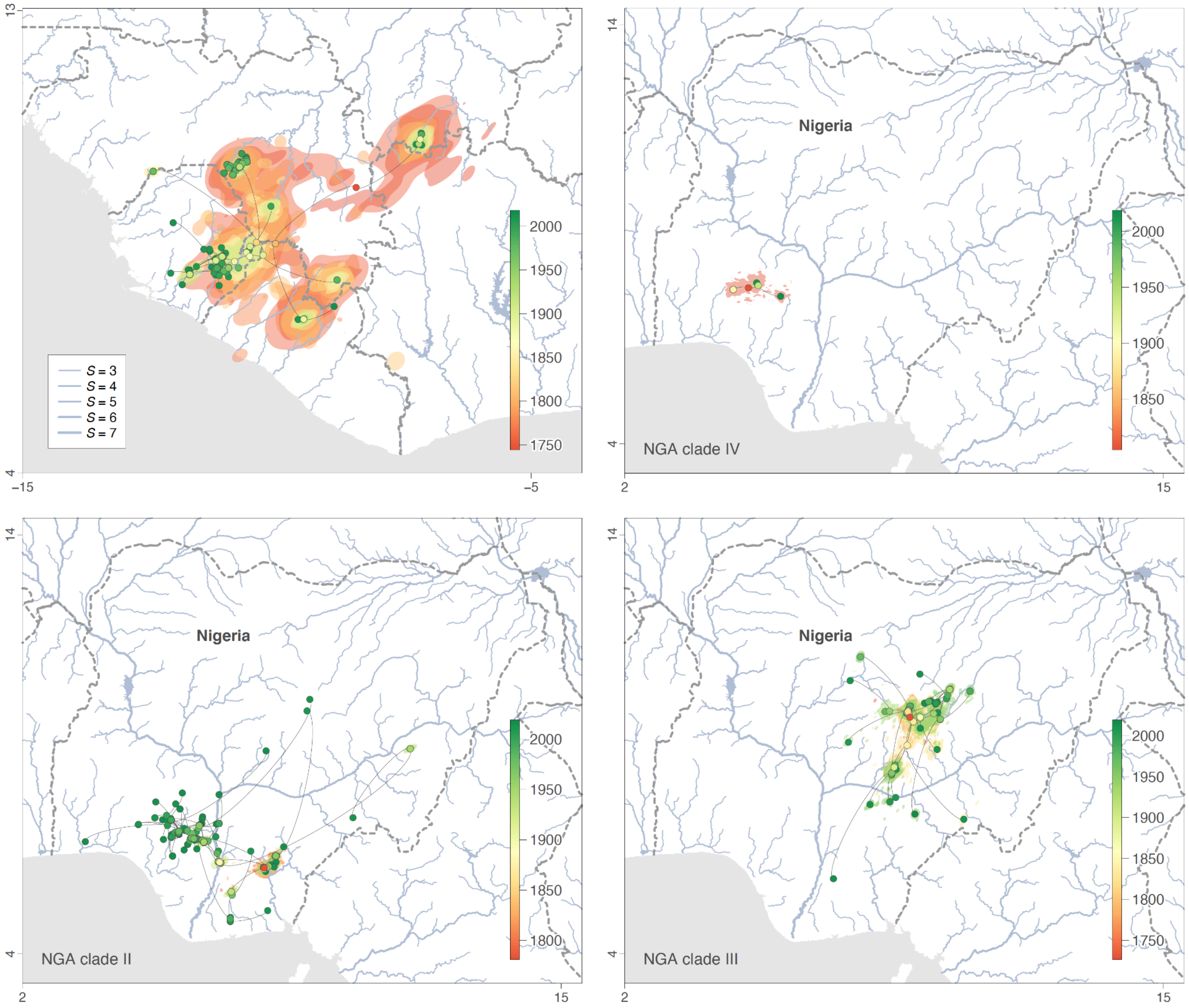


**Figure S10. Spatiotemporal diffusion of Lassa virus lineages in western Africa region and Nigeria, based on the analysis of segment S.** MCC (maximum clade credibility) tree obtained by continuous phylogeographic inference based on 1,000 posterior trees. A distinct phylogeographic analysis has been based on segments L and S as well as, in the case of the Nigerian data set, on clades I, II and III. These MCC trees are superimposed on 80% HPD reflecting phylogeographic uncertainty. Nodes of the trees, as well as HPD regions, are coloured from red (the time to the most recent common ancestor, TMRCA) to green (most recent sampling time), and oldest nodes (and corresponding HPD regions) are here plotted on top of youngest nodes. The trees are superimposed on maps displaying the main rivers present in the study area and classified according to their Strahler number *S*, which measures the importance of a river by counting the number of upstream rivers connected to it. International borders are represented by white dashed lines.

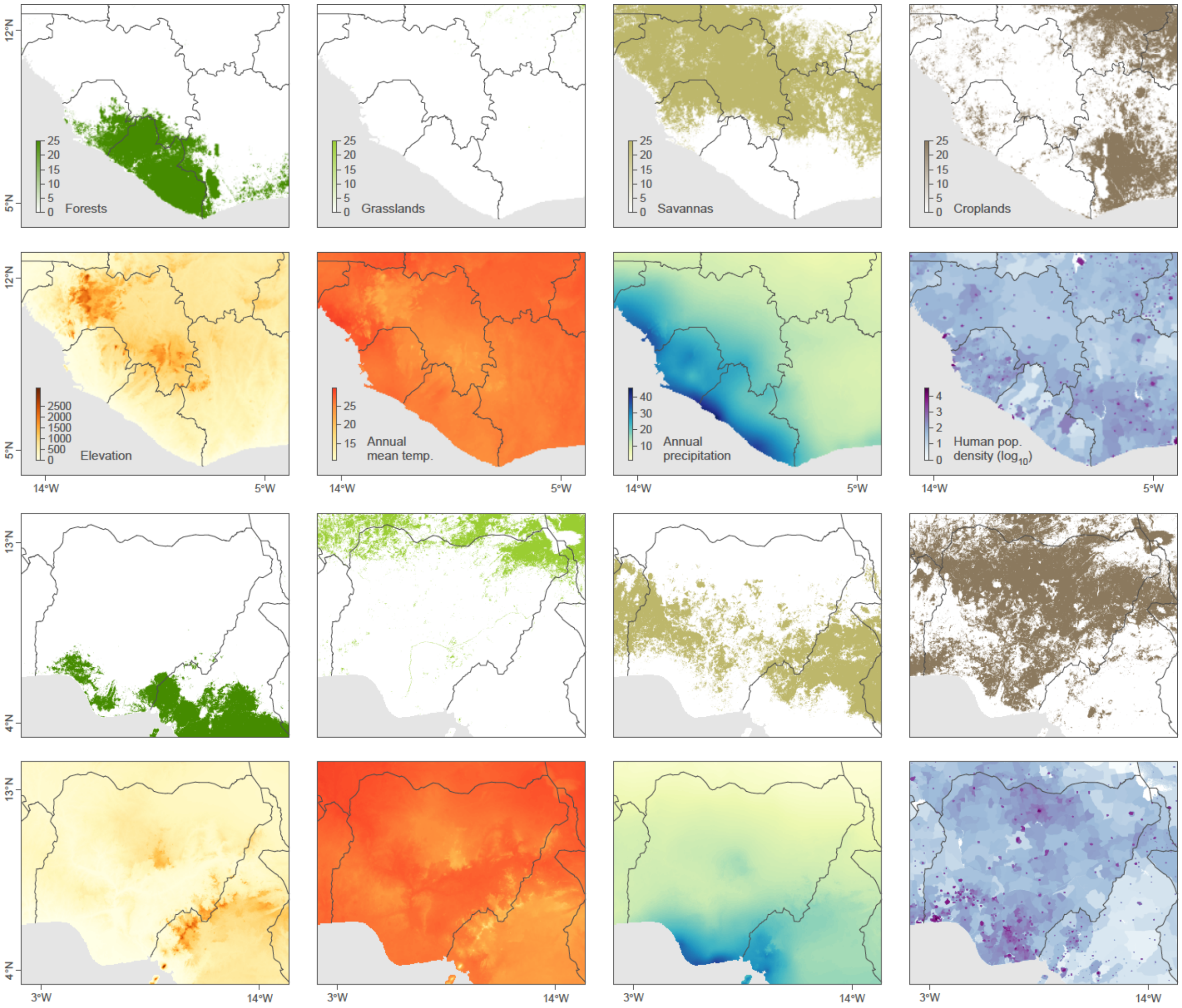


**Figure S11. Environmental variables tested for their impact on the dispersal velocity of Lassa virus lineages in western Africa and in Nigeria.** Elevation is reported in meters, mean annual temperature is reported in Celsius degrees, and annual precipitation is reported in meters per year.

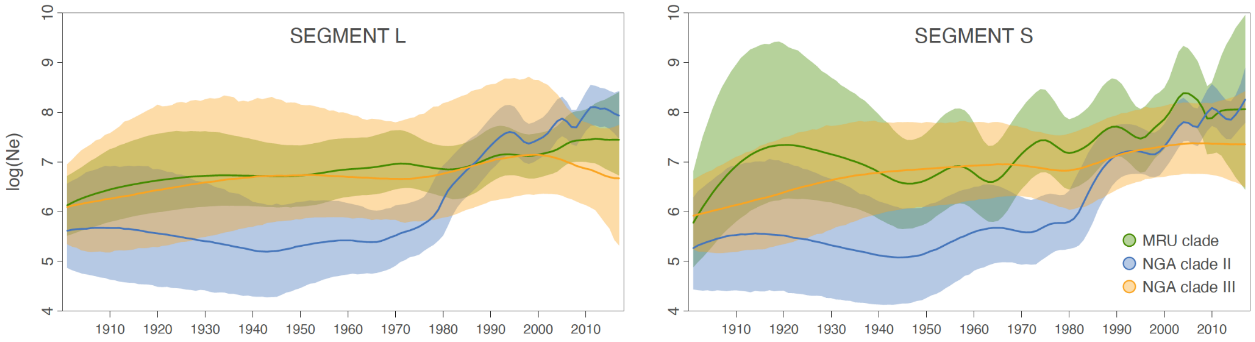


**Figure S12. Estimation of the viral effective population through time.** We here report Bayesian skygrid effective population size estimates based on the analysis of both segments (L and S) and for all main LASV clades. Solid curves and surrounding shaded polygons correspond to median estimates and 95% HPD region, respectively.

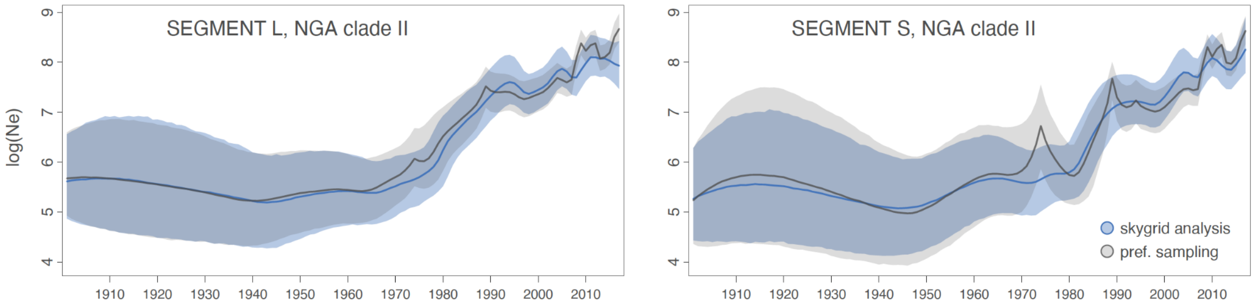


**Figure S13. Estimation of the viral effective population through time for NGA clade II.** We here report Bayesian skygrid effective population size estimates based on the analysis of both segments (L and S) for NGA clade II (in blue), as well as the results of the corresponding preferential sampling analysis (in grey). Contrary to the skygrid analysis, the preferential sampling analysis accounts for heterogeneous sampling density through time, which can improve estimates of global effective population size. In the graphs, solid curves and surrounding shaded polygons correspond to median estimates and 95% HPD region, respectively. See Figure S12 for a comparison with the skygrid reconstructions obtained for the MRU clade and NGA clade III.

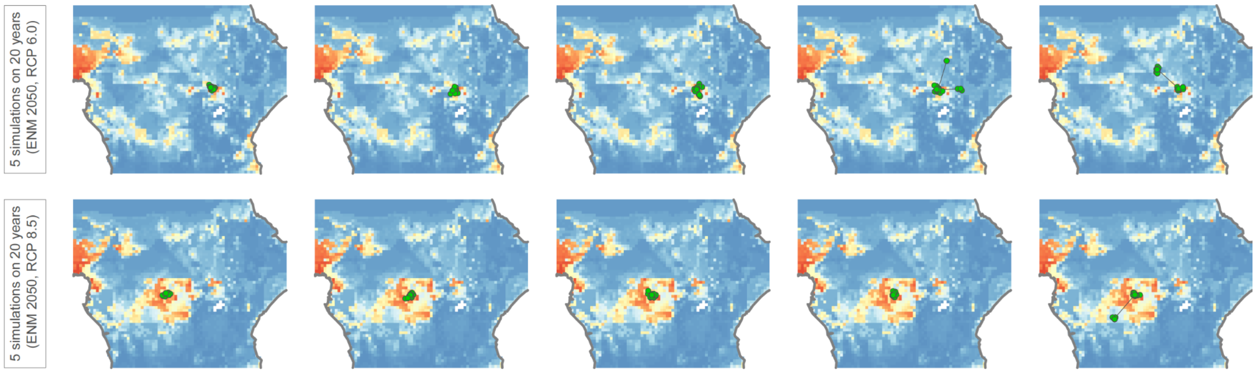


**Figure S14. Phylogeographic simulations of a viral spread following a successful introduction into a new ecologically suitable area.** Phylogeographic simulations are based on the tree topologies inferred for Lassa virus clade II (segment S). Five out of 1,000 simulated trees are shown superimposed on maps displaying future ecological suitability estimated for Lassa virus. All 1,000 trees were simulated starting from the same ancestral location and virus dispersal was constrained by ecological suitability using ecological niche projections for Lassa virus in 2050 according to scenarios RCP 6.0 (upper panels) and RCP 8.5 (lower panels).

**Table S1. Evolution of the population living in areas ecologically suitable for Lassa virus.** For each scenario (RCP 2.6, 6.0, and 8.5) and time period (2030, 2050, and 2070), we calculated the number of people at risk of exposure to Lassa virus (in millions) from index of human exposure (IHE) values ("projected"). To estimate the effect of the increase in population living in current suitable areas we combined current projected ecologically suitable areas with future population projections to re-estimate IHE values. To estimate the effect of population growth, we used current population projections combined with future projected ecologically suitable areas and re-estimated IHE values ("current pop."). For all population estimates, we provide the mean, and 95% HPD interval across estimates obtained with all climatic models and ecological niche model replicates. Counts for "endemic areas" correspond to the sum of the counts for Guinea, Liberia, Nigeria and Sierra Leone, counts for Africa are for the whole African continent. Here, values are only shown for the intermediate scenario, i.e. RCP 6.0. We also calculated the future population living in areas in 2070 by subtracting the future population living throughout all areas suitable in 2070 with the future population living in current suitable areas (not shown in table), obtaining the following estimates: 337 [260-405] million people.

| | $t_0$ | 2030 | | | 2050 | | | 2070 | | |
|---|---|---|---|---|---|---|---|---|---|---|
| | projected | projected | current area | current pop. | projected | current area | current pop. | projected | current area | current pop. |
| **Guinea** | 2 [2-2] | 8 [7-8] | 4 [4-5] | 4 [4-4] | 10 [9-11] | 5 [4-5] | 4 [4-5] | 12 [11-13] | 5 [5-6] | 5 [5-5] |
| **Liberia** | 1 [1-1] | 5 [4-5] | 3 [3-3] | 2 [2-2] | 7 [6-9] | 4 [4-4] | 2 [2-2] | 11 [10-12] | 5 [5-5] | 2 [2-2] |
| **Nigeria** | 76 [70-80] | 167 [154-172] | 173 [159-182] | 74 [69-76] | 255 [247-259] | 251 [231-265] | 79 [76-80] | 335 [328-341] | 321 [296-339] | 81 [79-82] |
| **Sierra Leone** | 3 [3-3] | 7 [7-7] | 7 [7-7] | 3 [3-3] | 9 [9-9] | 8 [8-9] | 3 [3-3] | 11 [11-11] | 10 [10-10] | 3 [3-3] |
| **Endemic areas** | 83 [76-87] | 187 [172-192] | 186 [172-196] | 83 [77-86] | 282 [271-288] | 268 [248-283] | 88 [85-90] | 368 [359-377] | 341 [315-359] | 91 [89-93] |
| **Africa** | 92 [83-98] | 253 [233-268] | 203 [186-214] | 113 [104-120] | 453 [414-497] | 288 [264-304] | 144 [128-157] | 700 [624-779] | 363 [333-384] | 179 [159-199] |

**Table S2. Impact of main rivers on the dispersal frequency of Lassa virus lineages.** The results are based on 100 posterior trees obtained by spatially-explicit phylogeographic inference. For each category of rivers, we report the total cost (*TC*, i.e. the sum of the least-cost values computed for each phylogenetic branch considered separately) and associated Bayes factor (BF) support indicating if *TC* is lower than expected by chance, i.e. lower than the value computed in the null dispersal model. Following Kass & Raftery (1995), we consider a BF value >20 as strong support for a significant correlation between the environmental distances and dispersal durations (in bold). A river is here characterised by its Strahler number (*S*), which measures its importance by looking at the number of upstream rivers connected to it. *k* is the rescaling parameter used to transform the initial raster and defines the additional resistance to movement when the cell does contain such a potential landscape barrier (see the Methods section for further detail). For each estimated *TC* value, we report both the median estimate and the 95% HPD interval, as well as the associated Bayes factor (BF) support. (*) Analyses based on main rivers associated with a *S* higher than five were only performed for lineages occurring in Nigeria.

| Rivers category | | Segment L | | Segment S | |
|---|---|---|---|---|---|
| | ***k*** | ***TC*** | **BF** | ***TC*** | **BF** |
| ***S* > 2** | 10 | 11692 [11315-12173] | 1.9 | 6930 [6356-7512] | 3.5 |
| | 100 | 15842 [14937-16890] | 1.6 | 9260 [8349-9800] | 5.2 |
| | 1000 | 32868 [27776-39707] | 1.8 | 19167 [15167-22463] | 3.3 |
| ***S* > 3** | 10 | 11258 [10881-11722] | 2.3 | 6674 [6124-7251] | 3.8 |
| | 100 | 13230 [12530-14010] | 2.6 | 7786 [7058-8352] | 5.7 |
| | 1000 | 27414 [23237-33115] | 1.0 | 16084 [12827-19439] | 1.9 |
| ***S* > 4** | 10 | 11076 [10699-11551] | 2.6 | 6573 [6032-7157] | 3.5 |
| | 100 | 11962 [11398-12618] | 4.3 | 6997 [6313-7652] | 5.7 |
| | 1000 | 18723 [16556-23418] | 4.0 | 10851 [8604-12816] | 7.3 |
| ***S* > 5*** | 10 | 106662 [103866-108759] | 0.7 | 57034 [55872-58355] | 0.7 |
| | 100 | 125682 [122927-128087] | 0.7 | 66919 [65547-68244] | 0.9 |
| | 1000 | 235351 [228537-242152] | 0.9 | 122884 [119306-125924] | 1.1 |
| ***S* > 6*** | 10 | 106076 [103352-108162] | 0.6 | 56758 [55570-58068] | 0.7 |
| | 100 | 122850 [120201-125005] | 0.4 | 65324 [64124-66637] | 0.5 |
| | 1000 | 193963 [189229-199856] | 0.3 | 101658 [99346-103853] | 0.4 |

**Table S3. Impact of several environmental factors on the dispersal velocity of Lassa virus tip branches, based on the analysis of segment L.** The results are based on 100 posterior trees obtained by spatially-explicit phylogeographic inference. For the main rivers *S* refers to the Strahler number, which measures the importance of a river by counting the number of upstream rivers connected to it. "C" and "R" indicate if the considered environmental raster was considered as a conductance ("C") or resistance factor ("R"), "*k*" is the rescaling parameter used to transform the initial raster, and "*S*" refers to the Strahler number that measures its importance by looking at the number of upstream rivers connected to it (see the text for further detail). For regression coefficients and *Q* values we report both the median estimate and the 95% HPD interval. The Bayes factor (BF) supports are only reported when p(*Q* > 0) is at least 90%. Following Kass & Raftery (1995), we consider a BF value >20 as strong support for a significant correlation between the environmental distances and dispersal durations (in bold). (*) Analyses based on main rivers associated with a Strahler number higher than 5 were only performed for lineages occurring in Nigeria.

| Path model | Environmental factor | *k* | Regression coefficient | *Q* statistic | p(*Q* > 0) | BF |
|---|---|---|---|---|---|---|
| **Least-cost algorithm** | forests (R) | 10 | 0.051 [0.011, 0.103] | -0.043 [-0.082, -0.013] | 0.01 | - |
| | | 100 | 0.006 [0.000, 0.043] | -0.080 [-0.144, -0.036] | 0.00 | - |
| | | 1000 | 0.002 [0.000, 0.036] | -0.083 [-0.146, -0.036] | 0.00 | - |
| | forests (C) | 10 | 0.091 [0.047, 0.140] | 0.002 [-0.025, 0.020] | 0.63 | - |
| | | 100 | 0.089 [0.049, 0.129] | -0.001 [-0.041, 0.023] | 0.47 | - |
| | | 1000 | 0.089 [0.049, 0.129] | -0.001 [-0.043, 0.025] | 0.47 | - |
| | grasslands (R) | 10 | 0.091 [0.040, 0.152] | 0.001 [-0.001, 0.002] | 0.71 | - |
| | | 100 | 0.084 [0.044, 0.136] | -0.006 [-0.024, 0.008] | 0.17 | - |
| | | 1000 | 0.020 [0.010, 0.049] | -0.069 [-0.127, -0.010] | 0.01 | - |
| | grasslands (C) | 10 | 0.094 [0.036, 0.169] | 0.002 [-0.009, 0.014] | 0.63 | - |
| | | 100 | 0.092 [0.028, 0.181] | 0.001 [-0.019, 0.027] | 0.52 | - |
| | | 1000 | 0.088 [0.024, 0.176] | -0.004 [-0.023, 0.025] | 0.37 | - |
| | savannas (R) | 10 | 0.148 [0.040, 0.264] | 0.052 [-0.005, 0.116] | 0.95 | 11.5 |
| | | 100 | 0.134 [0.033, 0.226] | 0.040 [-0.018, 0.093] | 0.93 | 5.2 |
| | | 1000 | 0.130 [0.029, 0.213] | 0.030 [-0.024, 0.091] | 0.86 | - |
| | savannas (C) | 10 | 0.064 [0.026, 0.115] | -0.024 [-0.080, 0.003] | 0.08 | - |
| | | 100 | 0.040 [0.016, 0.109] | -0.047 [-0.118, 0.000] | 0.03 | - |
| | | 1000 | 0.019 [0.007, 0.084] | -0.065 [-0.133, -0.007] | 0.02 | - |
| | croplands (R) | 10 | 0.093 [0.049, 0.143] | 0.001 [-0.044, 0.038] | 0.52 | - |
| | | 100 | 0.086 [0.045, 0.134] | -0.003 [-0.071, 0.044] | 0.45 | - |
| | | 1000 | 0.084 [0.044, 0.131] | -0.005 [-0.076, 0.044] | 0.43 | - |
| | croplands (C) | 10 | 0.114 [0.031, 0.183] | 0.016 [-0.020, 0.054] | 0.71 | - |
| | | 100 | 0.081 [0.009, 0.109] | -0.021 [-0.074, 0.032] | 0.20 | - |
| | | 1000 | 0.070 [0.001, 0.096] | -0.034 [-0.089, 0.026] | 0.11 | - |
| | elevation (R) | 10 | 0.145 [0.065, 0.230] | 0.055 [0.019, 0.091] | 1.00 | 19.0 |
| | | 100 | 0.168 [0.076, 0.269] | 0.074 [0.016, 0.136] | 1.00 | 10.1 |
| | | 1000 | 0.168 [0.075, 0.272] | 0.072 [0.014, 0.132] | 0.98 | 9.0 |
| | elevation (C) | 10 | 0.054 [0.025, 0.096] | -0.037 [-0.062, -0.010] | 0.00 | - |
| | | 100 | 0.040 [0.020, 0.070] | -0.052 [-0.089, -0.016] | 0.00 | - |
| | | 1000 | 0.036 [0.018, 0.064] | -0.055 [-0.094, -0.018] | 0.00 | - |
| | annual mean temperature (R) | 10 | 0.088 [0.037, 0.148] | -0.003 [-0.007, 0.000] | 0.01 | - |
| | | 100 | 0.088 [0.037, 0.147] | -0.003 [-0.007, 0.000] | 0.01 | - |
| | | 1000 | 0.088 [0.037, 0.147] | -0.003 [-0.007, 0.000] | 0.01 | - |
| | annual mean temperature (C) | 10 | 0.095 [0.041, 0.161] | 0.004 [0.001, 0.008] | 1.00 | 4.3 |
| | | 100 | 0.095 [0.041, 0.161] | 0.004 [0.001, 0.009] | 1.00 | 4.3 |
| | | 1000 | 0.095 [0.041, 0.162] | 0.004 [0.001, 0.009] | 1.00 | 4.3 |
| | annual precipitation (R) | 10 | 0.091 [0.033, 0.150] | -0.003 [-0.019, 0.006] | 0.27 | - |
| | | 100 | 0.089 [0.030, 0.148] | -0.004 [-0.025, 0.006] | 0.22 | - |
| | | 1000 | 0.089 [0.030, 0.148] | -0.005 [-0.025, 0.007] | 0.22 | - |
| | annual precipitation (C) | 10 | 0.088 [0.039, 0.154] | -0.002 [-0.009, 0.014] | 0.38 | - |
| | | 100 | 0.086 [0.039, 0.153] | -0.003 [-0.012, 0.017] | 0.36 | - |
| | | 1000 | 0.086 [0.039, 0.152] | -0.003 [-0.012, 0.017] | 0.36 | - |
| | human population density ($log_{10}$, R) | 10 | 0.083 [0.035, 0.152] | -0.006 [-0.016, 0.002] | 0.09 | - |
| | | 100 | 0.082 [0.035, 0.150] | -0.008 [-0.019, 0.002] | 0.07 | - |
| | | 1000 | 0.082 [0.035, 0.150] | -0.008 [-0.020, 0.002] | 0.06 | - |
| | human population density ($log_{10}$, C) | 10 | 0.097 [0.042, 0.149] | 0.002 [-0.008, 0.010] | 0.68 | - |
| | | 100 | 0.098 [0.043, 0.150] | 0.003 [-0.009, 0.012] | 0.71 | - |
| | | 1000 | 0.098 [0.043, 0.150] | 0.003 [-0.009, 0.012] | 0.70 | - |

| | | | | | | |
|---|---|---|---|---|---|---|
| | main rivers ($S$ > 2, R) | 10 | 0.066 [0.039, 0.117] | -0.002 [-0.006, 0.000] | 0.05 | - |
| | | 100 | 0.053 [0.032, 0.101] | -0.013 [-0.032, -0.002] | 0.01 | - |
| | | 1000 | 0.022 [0.012, 0.051] | -0.043 [-0.081, -0.015] | 0.00 | - |
| | main rivers ($S$ > 3, R) | 10 | 0.066 [0.040, 0.118] | -0.001 [-0.002, 0.001] | 0.20 | - |
| | | 100 | 0.055 [0.033, 0.106] | -0.010 [-0.022, -0.001] | 0.02 | - |
| | | 1000 | 0.019 [0.010, 0.045] | -0.047 [-0.092, -0.022] | 0.00 | - |
| | main rivers ($S$ > 4, R) | 10 | 0.067 [0.040, 0.119] | 0.000 [-0.001, 0.001] | 0.49 | - |
| | | 100 | 0.065 [0.038, 0.114] | -0.002 [-0.010, 0.004] | 0.26 | - |
| | | 1000 | 0.031 [0.015, 0.062] | -0.034 [-0.076, -0.015] | 0.00 | - |
| | main rivers ($S$ > 5, R) | 10 | 0.067 [0.040, 0.119] | 0.000 [-0.001, 0.001] | 0.48 | - |
| | | 100 | 0.065 [0.038, 0.114] | -0.002 [-0.010, 0.004] | 0.26 | - |
| | | 1000 | 0.031 [0.015, 0.062] | -0.034 [-0.076, -0.015] | 0.00 | - |
| | main rivers ($S$ > 6, R) | 10 | 0.066 [0.040, 0.119] | 0.000 [-0.001, 0.000] | 0.02 | - |
| | | 100 | 0.062 [0.035, 0.114] | -0.003 [-0.01, -0.001] | 0.01 | - |
| | | 1000 | 0.021 [0.008, 0.043] | -0.045 [-0.079, -0.027] | 0.00 | - |
| **Circuitscape algorithm** | forests (R) | 10 | 0.008 [0.001, 0.022] | -0.071 [-0.098, -0.037] | 0.00 | - |
| | | 100 | 0.001 [0.000, 0.009] | -0.079 [-0.107, -0.038] | 0.00 | - |
| | | 1000 | 0.001 [0.000, 0.008] | -0.080 [-0.108, -0.038] | 0.00 | - |
| | forests (C) | 10 | 0.093 [0.055, 0.127] | 0.015 [-0.004, 0.030] | 0.95 | 1.9 |
| | | 100 | 0.083 [0.050, 0.117] | 0.004 [-0.017, 0.026] | 0.73 | - |
| | | 1000 | 0.077 [0.047, 0.113] | -0.002 [-0.024, 0.025] | 0.44 | - |
| | grasslands (R) | 10 | 0.077 [0.043, 0.107] | -0.001 [-0.010, 0.007] | 0.42 | - |
| | | 100 | 0.031 [0.018, 0.059] | -0.045 [-0.073, -0.009] | 0.01 | - |
| | | 1000 | 0.011 [0.004, 0.028] | -0.067 [-0.097, -0.022] | 0.00 | - |
| | grasslands (C) | 10 | 0.077 [0.034, 0.103] | -0.004 [-0.008, 0.000] | 0.06 | - |
| | | 100 | 0.070 [0.028, 0.097] | -0.011 [-0.020, -0.003] | 0.00 | - |
| | | 1000 | 0.063 [0.024, 0.093] | -0.016 [-0.029, -0.006] | 0.00 | - |
| | savannas (R) | 10 | 0.127 [0.057, 0.188] | 0.050 [0.009, 0.095] | 0.99 | 13.3 |
| | | 100 | 0.121 [0.048, 0.187] | 0.038 [-0.008, 0.096] | 0.94 | **24.0** |
| | | 1000 | 0.116 [0.046, 0.184] | 0.034 [-0.012, 0.093] | 0.90 | 19.0 |
| | savannas (C) | 10 | 0.039 [0.017, 0.058] | -0.043 [-0.066, -0.016] | 0.00 | - |
| | | 100 | 0.028 [0.012, 0.051] | -0.050 [-0.078, -0.018] | 0.00 | - |
| | | 1000 | 0.028 [0.010, 0.057] | -0.051 [-0.083, -0.011] | 0.00 | - |
| | croplands (R) | 10 | 0.060 [0.031, 0.084] | -0.019 [-0.038, 0.004] | 0.08 | - |
| | | 100 | 0.054 [0.028, 0.079] | -0.025 [-0.047, 0.004] | 0.05 | - |
| | | 1000 | 0.053 [0.027, 0.078] | -0.026 [-0.049, 0.004] | 0.05 | - |
| | croplands (C) | 10 | 0.070 [0.021, 0.105] | -0.010 [-0.037, 0.024] | 0.20 | - |
| | | 100 | 0.046 [0.001, 0.072] | -0.037 [-0.071, 0.009] | 0.05 | - |
| | | 1000 | 0.042 [0.001, 0.064] | -0.040 [-0.074, -0.002] | 0.03 | - |
| | elevation (R) | 10 | 0.132 [0.064, 0.179] | 0.050 [0.024, 0.081] | 1.00 | 10.1 |
| | | 100 | 0.149 [0.077, 0.207] | 0.066 [0.027, 0.114] | 1.00 | 10.1 |
| | | 1000 | 0.150 [0.079, 0.210] | 0.067 [0.026, 0.118] | 1.00 | 11.5 |
| | elevation (C) | 10 | 0.039 [0.017, 0.060] | -0.040 [-0.057, -0.019] | 0.00 | - |
| | | 100 | 0.015 [0.006, 0.029] | -0.063 [-0.085, -0.031] | 0.00 | - |
| | | 1000 | 0.010 [0.004, 0.021] | -0.068 [-0.091, -0.034] | 0.00 | - |
| | annual mean temperature (R) | 10 | 0.076 [0.036, 0.106] | -0.004 [-0.008, -0.001] | 0.00 | - |
| | | 100 | 0.076 [0.036, 0.105] | -0.004 [-0.008, -0.002] | 0.00 | - |
| | | 1000 | 0.076 [0.036, 0.105] | -0.004 [-0.008, -0.002] | 0.00 | - |
| | annual mean temperature (C) | 10 | 0.083 [0.039, 0.112] | 0.003 [0.000, 0.007] | 0.99 | 0.8 |
| | | 100 | 0.084 [0.039, 0.112] | 0.003 [0.000, 0.008] | 0.99 | 0.8 |
| | | 1000 | 0.084 [0.039, 0.112] | 0.003 [0.000, 0.008] | 0.99 | 0.8 |
| | annual precipitation (R) | 10 | 0.050 [0.022, 0.071] | -0.030 [-0.040, -0.016] | 0.00 | - |
| | | 100 | 0.044 [0.018, 0.064] | -0.036 [-0.049, -0.019] | 0.00 | - |
| | | 1000 | 0.043 [0.018, 0.063] | -0.037 [-0.050, -0.020] | 0.00 | - |
| | annual precipitation (C) | 10 | 0.099 [0.049, 0.130] | 0.018 [0.009, 0.028] | 1.00 | 1.9 |
| | | 100 | 0.103 [0.052, 0.133] | 0.021 [0.010, 0.033] | 1.00 | 1.8 |
| | | 1000 | 0.103 [0.052, 0.134] | 0.021 [0.010, 0.033] | 1.00 | 1.8 |
| | human population density ($log_{10}$, R) | 10 | 0.066 [0.030, 0.094] | -0.012 [-0.019, -0.003] | 0.00 | - |
| | | 100 | 0.064 [0.029, 0.091] | -0.014 [-0.023, -0.004] | 0.00 | - |
| | | 1000 | 0.063 [0.028, 0.090] | -0.014 [-0.024, -0.004] | 0.00 | - |
| | human population | 10 | 0.093 [0.046, 0.122] | 0.009 [0.002, 0.018] | 0.99 | 13.3 |

| | | | | | | |
|---|---|---|---|---|---|---|
| | density ($log_{10}$, C) | 100 | 0.094 [0.046, 0.123] | 0.010 [0.001, 0.022] | 0.98 | 11.5 |
| | | 1000 | 0.094 [0.046, 0.123] | 0.010 [0.001, 0.022] | 0.98 | 11.5 |

**Table S4. Impact of several environmental factors on the dispersal velocity of Lassa virus tip branches, based on the analysis of segment S.** The results are based on 100 posterior trees obtained by spatially-explicit phylogeographic inference. For the main rivers *S* refers to the Strahler number, which measures the importance of a river by counting the number of upstream rivers connected to it. "C" and "R" indicate if the considered environmental raster was considered as a conductance ("C") or resistance factor ("R"), "*k*" is the rescaling parameter used to transform the initial raster, and "*S*" refers to the Strahler number that measures its importance by looking at the number of upstream rivers connected to it (see the text for further detail). For regression coefficients and *Q* values we report both the median estimate and the 95% HPD interval. The Bayes factor (BF) supports are only reported when p(*Q* > 0) is at least 90%. Following Kass & Raftery (1995), we consider a BF value >20 as strong support for a significant correlation between the environmental distances and dispersal durations (in bold). (*) Analyses based on main rivers associated with a Strahler number higher than 5 were only performed for lineages occurring in Nigeria.

| **Path model** | **Environmental factor** | ***k*** | **Regression coefficient** | ***Q* statistic** | **p(*Q* > 0)** | **BF** |
|---|---|---|---|---|---|---|
| **Least-cost algorithm** | forests (R) | 10 | 0.033 [0.017, 0.054] | -0.034 [-0.051, -0.018] | 0.00 | - |
| | | 100 | 0.002 [0.000, 0.010] | -0.063 [-0.085, -0.044] | 0.00 | - |
| | | 1000 | 0.001 [0.000, 0.006] | -0.065 [-0.086, -0.046] | 0.00 | - |
| | forests (C) | 10 | 0.083 [0.061, 0.112] | 0.018 [0.008, 0.027] | 1.00 | 1.9 |
| | | 100 | 0.087 [0.063, 0.118] | 0.021 [0.010, 0.035] | 1.00 | 1.3 |
| | | 1000 | 0.088 [0.065, 0.120] | 0.023 [0.011, 0.037] | 1.00 | 1.3 |
| | grasslands (R) | 10 | 0.068 [0.050, 0.088] | 0.001 [0.000, 0.003] | 0.97 | 5.7 |
| | | 100 | 0.068 [0.049, 0.095] | 0.002 [-0.007, 0.019] | 0.62 | - |
| | | 1000 | 0.030 [0.019, 0.077] | -0.035 [-0.056, 0.005] | 0.06 | - |
| | grasslands (C) | 10 | 0.068 [0.044, 0.093] | 0.001 [-0.007, 0.011] | 0.54 | - |
| | | 100 | 0.057 [0.034, 0.083] | -0.010 [-0.020, 0.006] | 0.10 | - |
| | | 1000 | 0.049 [0.028, 0.074] | -0.019 [-0.029, -0.001] | 0.02 | - |
| | savannas (R) | 10 | 0.078 [0.055, 0.098] | 0.010 [-0.007, 0.033] | 0.90 | 11.5 |
| | | 100 | 0.051 [0.030, 0.083] | -0.014 [-0.040, 0.026] | 0.11 | - |
| | | 1000 | 0.038 [0.017, 0.066] | -0.028 [-0.056, 0.013] | 0.04 | - |
| | savannas (C) | 10 | 0.078 [0.049, 0.111] | 0.011 [-0.004, 0.029] | 0.90 | 1.0 |
| | | 100 | 0.071 [0.044, 0.114] | 0.007 [-0.016, 0.040] | 0.67 | - |
| | | 1000 | 0.068 [0.044, 0.110] | 0.005 [-0.024, 0.039] | 0.57 | - |
| | croplands (R) | 10 | 0.109 [0.068, 0.147] | 0.040 [0.019, 0.061] | 1.00 | 1.9 |
| | | 100 | 0.123 [0.078, 0.168] | 0.053 [0.026, 0.082] | 1.00 | 2.0 |
| | | 1000 | 0.125 [0.079, 0.171] | 0.055 [0.026, 0.085] | 1.00 | 1.9 |
| | croplands (C) | 10 | 0.060 [0.040, 0.080] | -0.007 [-0.026, 0.010] | 0.16 | - |
| | | 100 | 0.017 [0.006, 0.034] | -0.048 [-0.072, -0.028] | 0.00 | - |
| | | 1000 | 0.003 [0.000, 0.012] | -0.062 [-0.085, -0.046] | 0.00 | - |
| | elevation (R) | 10 | 0.098 [0.066, 0.137] | 0.031 [0.015, 0.054] | 1.00 | 1.9 |
| | | 100 | 0.112 [0.069, 0.169] | 0.048 [0.020, 0.086] | 1.00 | 1.4 |
| | | 1000 | 0.113 [0.068, 0.172] | 0.050 [0.020, 0.090] | 1.00 | 1.4 |
| | elevation (C) | 10 | 0.046 [0.035, 0.058] | -0.020 [-0.032, -0.011] | 0.00 | - |
| | | 100 | 0.034 [0.025, 0.044] | -0.031 [-0.047, -0.018] | 0.00 | - |
| | | 1000 | 0.027 [0.019, 0.036] | -0.038 [-0.054, -0.024] | 0.00 | - |
| | annual mean temperature (R) | 10 | 0.065 [0.048, 0.084] | -0.001 [-0.004, 0.000] | 0.07 | - |
| | | 100 | 0.065 [0.048, 0.083] | -0.001 [-0.004, 0.000] | 0.07 | - |
| | | 1000 | 0.065 [0.048, 0.083] | -0.001 [-0.005, 0.000] | 0.07 | - |
| | annual mean temperature (C) | 10 | 0.068 [0.049, 0.089] | 0.002 [0.000, 0.004] | 0.98 | 0.4 |
| | | 100 | 0.068 [0.049, 0.089] | 0.002 [0.000, 0.005] | 0.98 | 0.4 |
| | | 1000 | 0.068 [0.049, 0.089] | 0.002 [0.000, 0.005] | 0.98 | 0.4 |
| | annual precipitation (R) | 10 | 0.056 [0.040, 0.070] | -0.010 [-0.017, -0.005] | 0.00 | - |
| | | 100 | 0.053 [0.038, 0.066] | -0.013 [-0.022, -0.007] | 0.00 | - |
| | | 1000 | 0.053 [0.038, 0.066] | -0.013 [-0.023, -0.007] | 0.00 | - |
| | annual precipitation (C) | 10 | 0.073 [0.052, 0.097] | 0.007 [0.003, 0.012] | 1.00 | 1.5 |
| | | 100 | 0.074 [0.053, 0.100] | 0.008 [0.004, 0.015] | 1.00 | 1.4 |
| | | 1000 | 0.074 [0.053, 0.100] | 0.008 [0.004, 0.015] | 1.00 | 1.4 |
| | human population density ($log_{10}$, R) | 10 | 0.064 [0.046, 0.085] | -0.002 [-0.005, 0.002] | 0.08 | - |
| | | 100 | 0.063 [0.046, 0.085] | -0.002 [-0.006, 0.002] | 0.09 | - |

| | | | | | | |
|---|---|---|---|---|---|---|
| | | 1000 | 0.063 [0.046, 0.085] | -0.002 [-0.006, 0.002] | 0.09 | - |
| | human population density ($log_{10}$, C) | 10 | 0.069 [0.051, 0.090] | 0.003 [0.000, 0.007] | 0.99 | 3.0 |
| | | 100 | 0.070 [0.052, 0.091] | 0.004 [0.000, 0.008] | 0.99 | 3.2 |
| | | 1000 | 0.070 [0.052, 0.091] | 0.004 [0.001, 0.008] | 0.99 | 3.2 |
| | main rivers ($S > 2$, R) | 10 | 0.077 [0.053, 0.114] | 0.000 [-0.002, 0.002] | 0.57 | - |
| | | 100 | 0.068 [0.047, 0.101] | -0.008 [-0.022, 0.002] | 0.06 | - |
| | | 1000 | 0.038 [0.026, 0.057] | -0.038 [-0.070, -0.017] | 0.00 | - |
| | main rivers ($S > 3$, R) | 10 | 0.076 [0.053, 0.112] | -0.001 [-0.002, 0.000] | 0.16 | - |
| | | 100 | 0.068 [0.048, 0.101] | -0.008 [-0.019, 0.001] | 0.04 | - |
| | | 1000 | 0.033 [0.021, 0.056] | -0.044 [-0.074, -0.021] | 0.00 | - |
| | main rivers ($S > 4$, R) | 10 | 0.076 [0.053, 0.113] | 0.000 [-0.001, 0.000] | 0.31 | - |
| | | 100 | 0.073 [0.050, 0.105] | -0.003 [-0.011, 0.002] | 0.10 | - |
| | | 1000 | 0.036 [0.024, 0.048] | -0.039 [-0.069, -0.020] | 0.00 | - |
| | main rivers ($S > 5$, R) | 10 | 0.076 [0.053, 0.113] | 0.000 [-0.001, 0.000] | 0.30 | - |
| | | 100 | 0.073 [0.050, 0.105] | -0.003 [-0.011, 0.001] | 0.09 | - |
| | | 1000 | 0.036 [0.024, 0.048] | -0.040 [-0.069, -0.020] | 0.00 | - |
| | main rivers ($S > 6$, R) | 10 | 0.076 [0.052, 0.112] | -0.001 [-0.001, 0.000] | 0.02 | - |
| | | 100 | 0.069 [0.046, 0.102] | -0.007 [-0.013, -0.003] | 0.01 | - |
| | | 1000 | 0.020 [0.011, 0.031] | -0.057 [-0.089, -0.037] | 0.00 | - |
| **Circuitscape algorithm** | forests (R) | 10 | 0.008 [0.001, 0.016] | -0.068 [-0.085, -0.050] | 0.00 | - |
| | | 100 | 0.001 [0.000, 0.004] | -0.075 [-0.096, -0.056] | 0.00 | - |
| | | 1000 | 0.001 [0.000, 0.003] | -0.075 [-0.097, -0.056] | 0.00 | - |
| | forests (C) | 10 | 0.103 [0.079, 0.127] | 0.027 [0.018, 0.042] | 1.00 | 3.5 |
| | | 100 | 0.114 [0.089, 0.143] | 0.037 [0.027, 0.057] | 1.00 | 4.3 |
| | | 1000 | 0.119 [0.094, 0.151] | 0.043 [0.033, 0.065] | 1.00 | 5.2 |
| | grasslands (R) | 10 | 0.077 [0.058, 0.102] | 0.000 [-0.005, 0.015] | 0.54 | - |
| | | 100 | 0.039 [0.024, 0.083] | -0.036 [-0.059, 0.004] | 0.05 | - |
| | | 1000 | 0.018 [0.008, 0.060] | -0.056 [-0.083, -0.019] | 0.01 | - |
| | grasslands (C) | 10 | 0.070 [0.053, 0.093] | -0.006 [-0.013, -0.002] | 0.00 | - |
| | | 100 | 0.056 [0.038, 0.081] | -0.019 [-0.030, -0.010] | 0.00 | - |
| | | 1000 | 0.046 [0.029, 0.071] | -0.028 [-0.042, -0.019] | 0.00 | - |
| | savannas (R) | 10 | 0.042 [0.028, 0.063] | -0.034 [-0.048, -0.013] | 0.00 | - |
| | | 100 | 0.020 [0.011, 0.037] | -0.055 [-0.074, -0.031] | 0.00 | - |
| | | 1000 | 0.016 [0.009, 0.032] | -0.059 [-0.078, -0.035] | 0.00 | - |
| | savannas (C) | 10 | 0.060 [0.039, 0.078] | -0.017 [-0.031, -0.007] | 0.00 | - |
| | | 100 | 0.055 [0.034, 0.078] | -0.020 [-0.044, -0.001] | 0.03 | - |
| | | 1000 | 0.058 [0.033, 0.089] | -0.015 [-0.048, 0.011] | 0.16 | - |
| | croplands (R) | 10 | 0.092 [0.062, 0.116] | 0.016 [0.002, 0.027] | 0.98 | 0.8 |
| | | 100 | 0.092 [0.060, 0.117] | 0.016 [0.000, 0.030] | 0.97 | 1.3 |
| | | 1000 | 0.092 [0.060, 0.118] | 0.016 [0.000, 0.030] | 0.95 | 1.3 |
| | croplands (C) | 10 | 0.018 [0.007, 0.036] | -0.057 [-0.073, -0.041] | 0.00 | - |
| | | 100 | 0.001 [0.000, 0.008] | -0.075 [-0.096, -0.055] | 0.00 | - |
| | | 1000 | 0.000 [0.000, 0.004] | -0.075 [-0.098, -0.057] | 0.00 | - |
| | elevation (R) | 10 | 0.112 [0.082, 0.148] | 0.036 [0.016, 0.059] | 1.00 | 1.0 |
| | | 100 | 0.115 [0.075, 0.161] | 0.037 [0.010, 0.070] | 1.00 | 1.1 |
| | | 1000 | 0.114 [0.073, 0.161] | 0.036 [0.008, 0.070] | 1.00 | 1.2 |
| | elevation (C) | 10 | 0.040 [0.028, 0.054] | -0.035 [-0.049, -0.025] | 0.00 | - |
| | | 100 | 0.015 [0.009, 0.023] | -0.061 [-0.081, -0.045] | 0.00 | - |
| | | 1000 | 0.006 [0.003, 0.011] | -0.070 [-0.093, -0.053] | 0.00 | - |
| | annual mean temperature (R) | 10 | 0.074 [0.056, 0.097] | -0.001 [-0.005, 0.001] | 0.09 | - |
| | | 100 | 0.074 [0.056, 0.097] | -0.002 [-0.005, 0.001] | 0.07 | - |
| | | 1000 | 0.074 [0.056, 0.097] | -0.002 [-0.006, 0.001] | 0.07 | - |
| | annual mean temperature (C) | 10 | 0.079 [0.059, 0.103] | 0.003 [-0.001, 0.006] | 0.96 | 0.2 |
| | | 100 | 0.079 [0.059, 0.104] | 0.004 [0.000, 0.007] | 0.97 | 0.2 |
| | | 1000 | 0.080 [0.059, 0.104] | 0.004 [0.000, 0.007] | 0.97 | 0.2 |
| | annual precipitation (R) | 10 | 0.045 [0.030, 0.063] | -0.030 [-0.038, -0.023] | 0.00 | - |
| | | 100 | 0.039 [0.025, 0.055] | -0.037 [-0.046, -0.028] | 0.00 | - |
| | | 1000 | 0.038 [0.024, 0.054] | -0.038 [-0.047, -0.029] | 0.00 | - |
| | annual precipitation (C) | 10 | 0.100 [0.075, 0.126] | 0.023 [0.017, 0.032] | 1.00 | 2.3 |
| | | 100 | 0.103 [0.079, 0.129] | 0.027 [0.018, 0.037] | 1.00 | 2.3 |
| | | 1000 | 0.104 [0.079, 0.130] | 0.027 [0.018, 0.038] | 1.00 | 2.3 |

| | | | | | | |
|---|---|---|---|---|---|---|
| | human population density (log$_{10}$, R) | 10 | 0.064 [0.048, 0.086] | -0.011 [-0.016, -0.005] | 0.00 | - |
| | | 100 | 0.062 [0.046, 0.083] | -0.013 [-0.019, -0.006] | 0.00 | - |
| | | 1000 | 0.062 [0.046, 0.083] | -0.014 [-0.019, -0.006] | 0.00 | - |
| | human population density (log$_{10}$, C) | 10 | 0.083 [0.064, 0.108] | 0.007 [0.001, 0.012] | 1.00 | 11.5 |
| | | 100 | 0.083 [0.064, 0.107] | 0.007 [0.000, 0.013] | 0.98 | 6.7 |
| | | 1000 | 0.083 [0.064, 0.107] | 0.007 [0.000, 0.013] | 0.97 | 6.1 |

**Table S5. Online platforms used to retrieve sampling coordinates for Lassa virus samples.**

| Original dataset | URL |
|---|---|
| OpenStreetMap | https://data.humdata.org/dataset/open-street-map-data-on-guinea-liberia-and-sierra-leone |
| GeoNames gazetteers | https://www.geonames.org |
| UN-OCHA (for Sierra Leone) | https://data.humdata.org/dataset/sierra-leone-settlements |
| UN-OCHA (for Liberia) | https://data.humdata.org/dataset/liberia-settlements-0 |
| Who's on First gazetteer | https://www.whosonfirst.org/docs |
| Google geocoding API | https://developers.google.com/maps/documentation/geocoding/overview |

**Table S6. Source of data for each environmental raster.**

| Original raster | Source | URL |
|---|---|---|
| Elevation raster | SRTM (Shuttle Radar Topography Mission) | webmap.ornl.gov |
| Land cover raster | IGBP (International Geosphere Biosphere Programme) – categorical raster | www.igbp.net |
| Annual mean temperature | WorldClim database, version 2.0 (bioclimatic variable “bio1”) | worldclim.org |
| Annual Precipitation | WorldClim database, version 2.0 (bioclimatic variable “bio12”) | worldclim.org |
| Rivers of Africa | Food and Agriculture Organization of the United Nations | data.apps.fao.org |
